# Horizontal transfer of prokaryotic cytolethal distending toxin B genes to eukaryotes

**DOI:** 10.1101/544197

**Authors:** Kirsten I. Verster, Jennifer H. Wisecaver, Rebecca P. Duncan, Marianthi Karageorgi, Andrew D. Gloss, Ellie Armstrong, Donald K. Price, Aruna R. Menon, Zainab M. Ali, Noah K. Whiteman

## Abstract

Cytolethal distending toxins (CDTs) are tripartite eukaryotic genotoxins encoded in diverse bacterial and phage genomes. The cdtB subunit is a DNAse that causes eukaryotic cell cycle arrest and apoptosis, and in one context, is associated with resistance against parasitoid wasp infections. Here we report the discovery of functional *cdtB* copies in the nuclear genomes of insect species from two distantly related insect orders, including fruit flies (Diptera: Drosophilidae) and aphids (Hemiptera: Aphididae). Insect cdtB copies are most closely related to bacteriophage copies, were horizontally transferred to insect genomes > 40 million years ago and encode a protein that retains ancestral DNase activity. This phage-derived toxin has been domesticated by diverse insects and we hypothesize that it is used as a defensive weapon against parasitoid wasps.

**One Sentence Summary:** We report horizontal transfer of the gene *cytolethal distending toxin B*, which encodes a DNase, into eukaryotic genomes from bacteriophage.

**Significance:** Cytolethal distending toxins (CDTs) are secreted by diverse pathogenic bacterial species to kill animal cells. The cdtB subunit enters cell nuclei, damaging the DNA and leading to mitotic arrest and apoptosis. In the pea aphid, a bacterial endosymbiont provides protection against wasp attack, possibly via *cdtB.* We discovered that this same endosymbiont-encoded lineage of *cdtB* was transferred to the genomes of Diptera and Hemiptera species and retains ancestral DNase activity. This is the first report of *cdtB* outside of bacteria or phages. A toxin that first evolved to kill eukaryotic cells has been co-opted by insects, potentially to their benefit.

## Main Text

Cytolethal distending toxins (CDTs) are widespread intracellular-acting eukaryotic genotoxins encoded by a gene family restricted to Actinobacteria, Proteobacteria and bacteriophage genomes (*1*). CDTs are found in diverse pathogens, including *Campylobacter jejuni, Escherichia coli, Salmonella enterica*, and *Yersinia pestis* and may be a cause of irritable bowel syndrome (*1*). CDT holotoxin is an AB_2_ toxin typically encoded in a three-gene operon (*cdtA, cdtB,* and *cdtC*) (*2*) and cdtB is the catalytic subunit necessary for DNase activity (*3, 4*). CdtB nicking leads to DNA damage in eukaryotic cells followed by cell cycle arrest, cellular distention and death (*5*).

Although cdtB is a eukaryotic genotoxin, in one context it is associated with increased fitness of eukaryotes. Some strains of the bacterium *Candidatus* Hamiltonella defensa, a secondary endosymbiont of the pea aphid (*Acyrthosiphon pisum*), are infected with strains of the lysogenic bacteriophage APSE (*6, 7*). APSE-positive *Ca.* H defensa strains confer protection from attack by parasitoid braconid wasps that insert eggs into aphids (*8*). Comparative genomic studies point to *cdtB*, which is encoded in the genome of phage strain APSE-2, as a likely candidate underlying this protective effect (*6*–*8*).

We used a sequence similarity-based screen (*9*) to identify a *cdtB* homolog as a horizontal gene transfer (HGT) candidate in a *de novo* genome assembly of the drosophilid fly *Scaptomyza flava.* To identify *cdtB* copies in genomes of other eukaryotes, we executed TBLASTN (*10*) searches of the NBCI refseq database (which includes all eukaryotes), all NCBI ‘*Drosophila’* genomes, and the genomes of 11 unpublished Hawaiian *Drosophila* species. We found high-confidence hits to *cdtB* homologs in the drosophilid species *Dr. ananassae, Dr. bipectinata* (both in the ananassae subgroup) and *Dr. biarmipes,* the Hawaiian *Dr. primaeva,* and the aphid species *Myzus persicae* (**Table S1a)**. We also discovered *cdtB* orthologs in the transcriptomes of two other species in the ananassae subgroup, *Dr. pseudoananassae* and *Dr. ercepeae* (*11*). We subsequently searched all available AphidBase genomes and found single high-confidence hits to *cdtB* homologs in the Russian wheat aphid (*Diuraphis noxia*) and the black cherry aphid (*M. cerasi*), both in the Macrosiphini (**Table S1b**).

Putative HGT events can be due to microbial contamination arising from low-quality genome assemblies (*12*), so we used several methods to address these possibilities (*9*). First, *cdtB* was identified on scaffolds in species with high-quality genome assemblies (**Table S2)**. The presence of *cdtB* was verified by PCR and Sanger sequencing of both genomic and complementary DNA **(Table S3; Figure S1**). *CdtB*, when present, was found in all transcriptomes except that of *Di. noxia* (**Table S1**). The transcriptome libraries we searched were enriched for polyadenylated mRNA, suggesting insect *cdtB* was not due to bacterial contamination since bacteria typically lack 3’-polyA tails (*13*). Additionally, mRNA sequences of *cdtB* from all insect species (other than *S. flava*) contain at least three exons separated by intronic splice sites (*14*), which are rare in bacteria. The absence of *cdtB* transcripts in *Di. noxia,* coupled with a frame-shifting deletion and stop codon in the first (and only) predicted exon suggests that this *cdtB* fragment is a pseudogene in this species.

Phylogenetic conflict between gene tree and species tree topologies provides additional support for HGT (*15*). To evaluate this and determine the potential source of insect-encoded *cdtB*, we reconstructed a cdtB protein phylogeny using all available sequences (*9*). Viral, bacterial, and metazoan cdtB sequences were downloaded from the NCBI refseq protein database, aligned and used to create a protein tree (**Figure 1**, full phylogeny in **Figure S2**). The cdtB phylogeny reveals that all insect cdtB sequences form a clade with cdtB sequences from *Ca.* H. defensa and APSE-2. A HGT event from an APSE-2 ancestor to eukaryotes is further supported by the case of *Dr. bipectinata,* in which two *cdtB* copies are present in tandem array. One of the two *cdtB* copies in *Dr. bipectinata* is fused with a homolog of an unrelated AB toxin, *apoptosis inducing protein 56,* found immediately downstream of *cdtB* in *Ca.* H. defensa. This chimeric *ctdB+aip56* sequence is expressed as mRNA in *Dr. bipectinata*. Synteny between *Dr. bipectinata* and *Ca.* H. defensa suggests the two genes were horizontally transferred together (see **Supplementary Text**) from a bacterial or phage ancestor prior to the divergence of the extant ananassae spp. subgroup *Drosophila* species. This hypothesis is supported by the presence of homologous *cdtB+aip56* chimeric sequences in two other ananassae subgroup species, though it has been lost in *D. ananassae*.

**Fig. 1.**
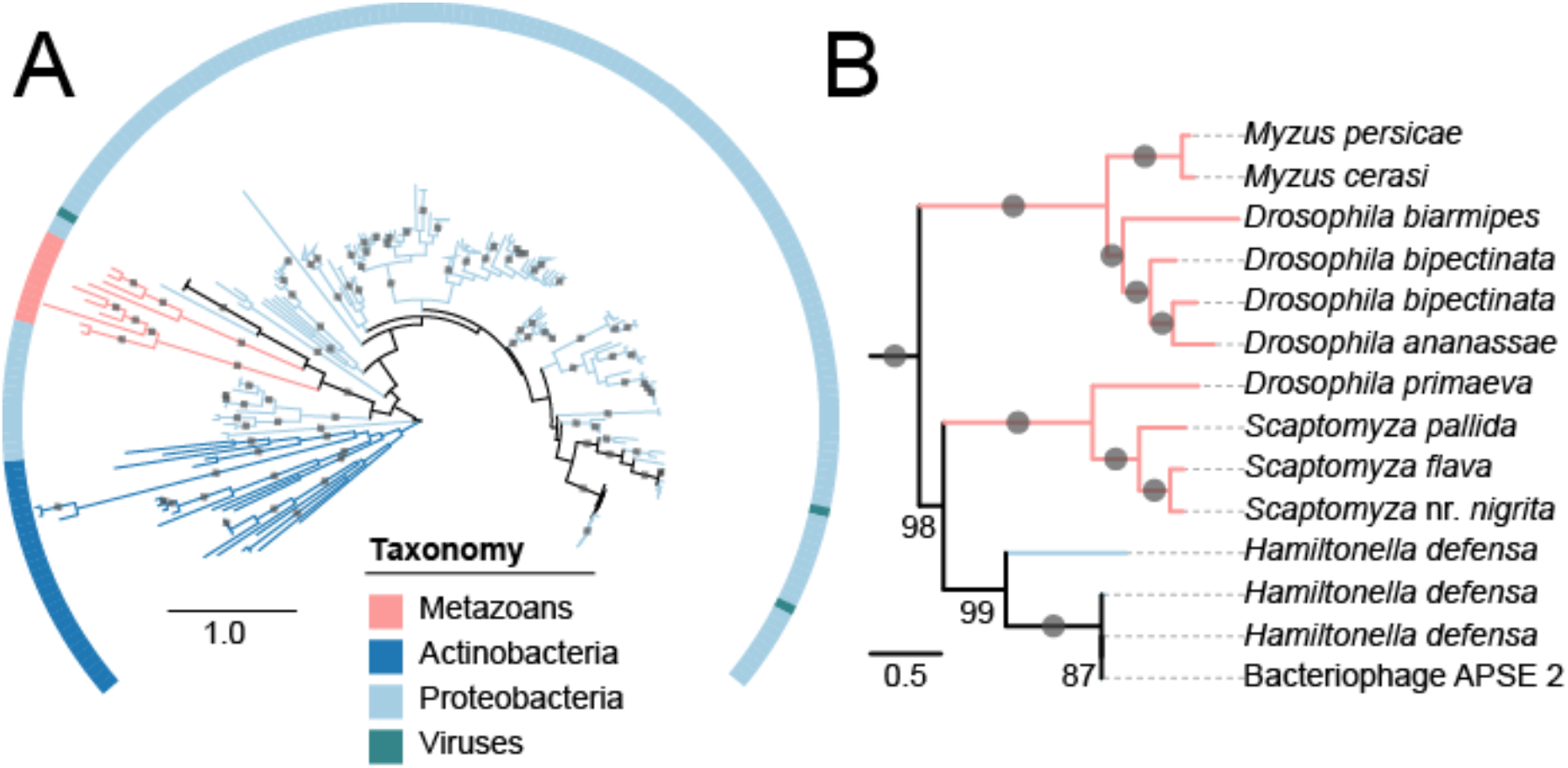
CdtB protein phylogeny indicates HGT into insects from bacteria. (A) ML phylogeny of cdtB from across the tree of life. Tree is midpoint rooted and branches with 100% bootstrap support are indicated by grey circles. Four clades consisting of highly similar sequences from Proteobacteria were collapsed for clarity. The full phylogeny is available in **Fig. S2.** (B) Detailed view of insect cdtB clades. Numbers below branches indicate percent bootstrap support when < 100.

Our data suggest two independent acquisitions of intron-bearing and intronless insect *cdtB.* The cdtB phylogeny resolves two insect-encoded sub-clades, one containing all cdtB sequences encoded by insect-encoded, intron-bearing *cdtB* copies (*Myzus* spp., *Dr. biarmipes,* and ananassae spp.) and the other containing all intron-less insect-encoded cdtB copies (*Scaptomyza* spp. + *D. primaeva*), which is in turn sister to the clade containing cdtB from *Ca.* H. defensa and APSE-2 genomes. Furthermore, an approximately unbiased test forcing monophyly of drosophilid cdtB is slightly worse (p=0.059) than the recovered cdtB phylogeny, suggesting that the intronless cdtB and the intron-bearing *cdtB* were independently transferred into insects.

In order to understand the number and timing of horizontal transfer of *cdtB* in insects, we reconstructed drosophilid and aphid species phylogenies and mapped *cdtB* evolution on these trees (*9*). We first constructed a drosophilid species phylogeny including all *Drosophila* and *Scaptomyza* genomes scanned for *cdtB*. We performed ancestral state reconstruction (ASR) for the origin of *cdtB* to estimate the number and timing of HGT events across the drosophilid species phylogeny. This analysis, coupled with a clear pattern of conserved synteny within clades, suggests that *cdtB* was acquired three times in drosophilids: (1) prior to the divergence of the ananassae subgroup (94% posterior clade probability, or PP) ca. 21 million years ago (mya) (*16*), (2) following the split between *Dr. biarmipes* and *Dr. suzukii* (98% PP) ca. 7.3 ± 2.5 mya (*17*), and (3) in an ancestor common to *S. flava* and *Dr. primaeva* (13% PP) ca. 24 ± 7 mya (*18*) **(Figure 2A**). While the likelihood that *cdtB* was present in the common ancestor of *Dr. primaeva* and *S. flava* is low based on ASR, synteny suggests that a single HGT event occurred in a common ancestor of these two species. None of the genomes (out of 10 surveyed) from the more recently derived Hawaiian *Drosophila* species sister to *Dr. primaeva* were found to encode a *cdtB* copy. Thus, *cdtB* was most likely lost prior to the divergence of the picture wing clade, ca. 7 ± 4 mya (*18*). We did not perform ASR in aphids due to limited availability of sequenced aphid genomes. However, *cdtB* was syntenic in *Di. noxia, M. cerasi* and *M. persicae,* distantly related members of the Macrosiphini. We hypothesize that *cdtB* was horizontally transferred into a common ancestor of these three aphid species (41 ± 5 mya (*19*)). While a functional copy was retained in *M. persicae* and *M. cerasi,* it was pseudogenized in *Di. noxia* and lost completely in *A. pisum* **(Figure 2B**).

**Fig. 2.**
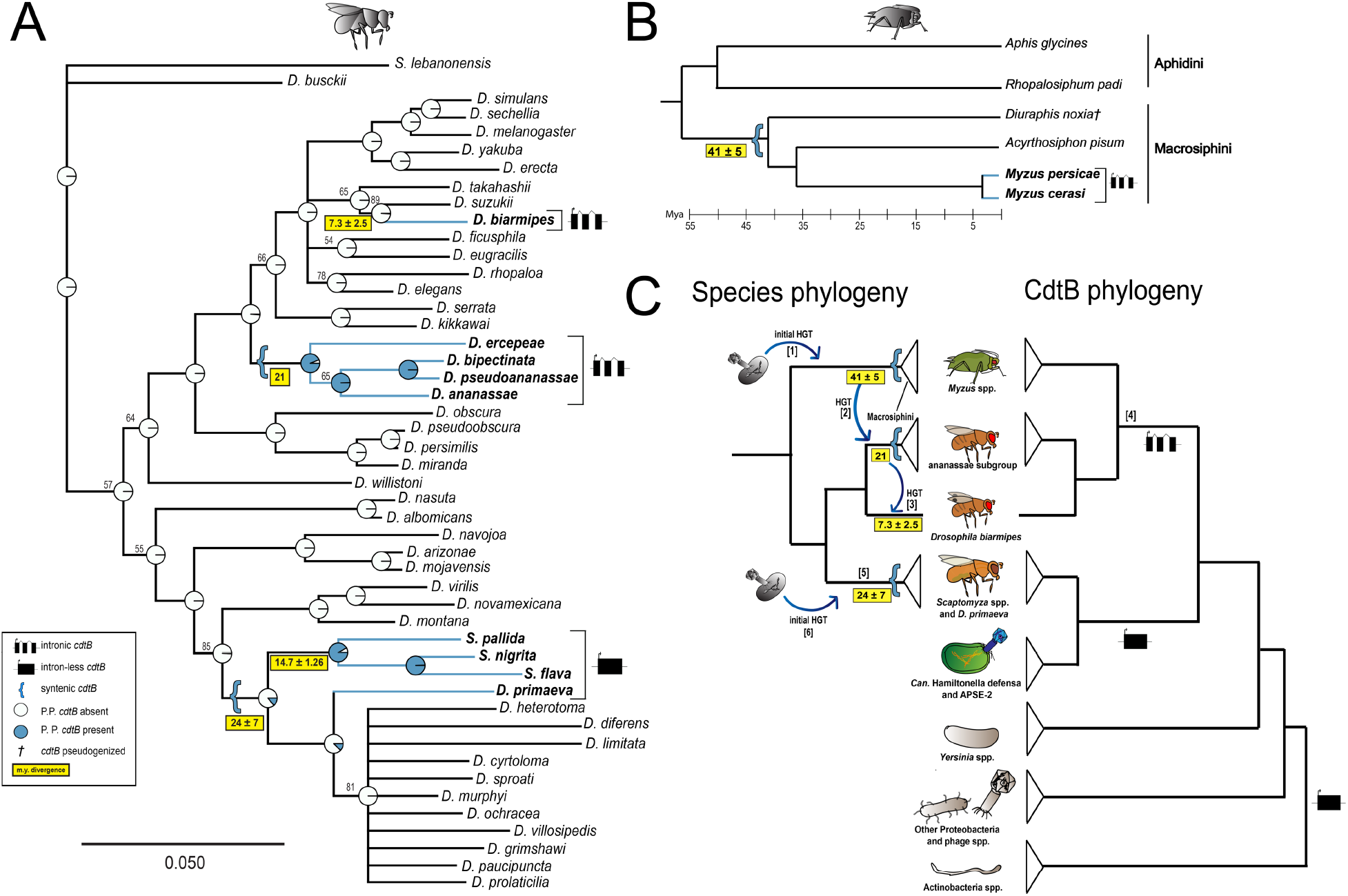
Species phylogenies show *cdtB* was transferred into, and possibly between, genomes of distant insect lineages. **A.** ML phylogeny of drosophilid species. Node labels indicate bootstraps if <90% or are collapsed to polytomies if <50%. ASR shows posterior probability (P.P.) of *cdtB* at nodes. **B.** Phylogeny of Aphidinae species. Branch lengths drawn approximately to scale using divergence dates from (*19, 30*). **C.** Simplified paired cdtB and species phylogenies. Blue arrows suggest possible HGT directions and bracketed numbers are described here. Possible initial prokaryote-eukaryote HGTs are [1,6]. We hypothesize an initial HGT of *cdtB* from bacteria or phage integrated into an aphid nuclear genome [1] and was lost or pseudogenized in some aphid lineages (**2B**). We then posit an inter-ordinal transfer [2] from a *Myzus* spp. ancestor to an ananassae subgroup spp. ancestor, followed by inter-specific transfer [3] to a *D. biarmipes* ancestor. This transfer sequence is supported by subclade ages, conserved intron splice sites in [4], and the regional co-occurrence of these subclades (*31, 32*). However, conserved exon structure in [4] could also arise from convergence. CdtB in [5] could have evolved independently, or was derived from the same HTG as [4] but failed to acquire introns.

Interestingly, *cdtB* copies with three exons (*Myzus* spp., *Dr. biarmipes,* and ananassae spp.) share identical splice junctions (**Figure S3**), which indicates either convergent origins of a modular exonic structure or that intron-bearing *cdtB* copies share a common ancestor and have been transferred horizontally between these distantly related insect lineages after an initial HGT event into one insect (*20*). HGT within eukaryotes could be mediated by several mechanisms, including predaceous mites (*21*), bracovirus (by parasitoid wasps intermediaries) and helitrons (*22*). We illustrate hypotheses on the order and timing of *cdtB* HGT in **Figure 2C**.

There are many examples of genes derived from prokaryote-to-eukaryote HGT events stably integrating into nuclear genomes, and this process often involves optimizing the transferred genes for expression in eukaryotic cells (*23*). All insect-encoded *cdtB* copies exhibit features common to eukaryotic transcription initiation and termination (**Figure S4**; **Supplementary Text**). Additionally, insect *cdtB* copies have polyadenylated mRNA, 5’ and 3’ untranslated regions, and introns (except for *Scaptomyza* spp. + *Dr. primaeva*), which may modulate eukaryotic transcription/translation (*13, 24, 25*).

Expression patterns of HTGs often evolve to become finely tuned to eukaryotic cellular environments (*24, 26*). We evaluated if *cdtB* shows differential expression patterns throughout development in two drosophilids that represent species with intron-bearing and intronless *cdtB* (*9*). Consistent with a potential role in parasitoid resistance, we predicted that *cdtB* expression would be highest in larvae, the developmental stage most prone to parasitoid attack in drosophilids (*27*). We used RT-qPCR in larvae, pupae, and adult males and females of *S. flava* and *Dr. ananassae* and found that *cdtB* expression was indeed highest in larvae of both species **(Figure 3)**.

**Fig. 3.**
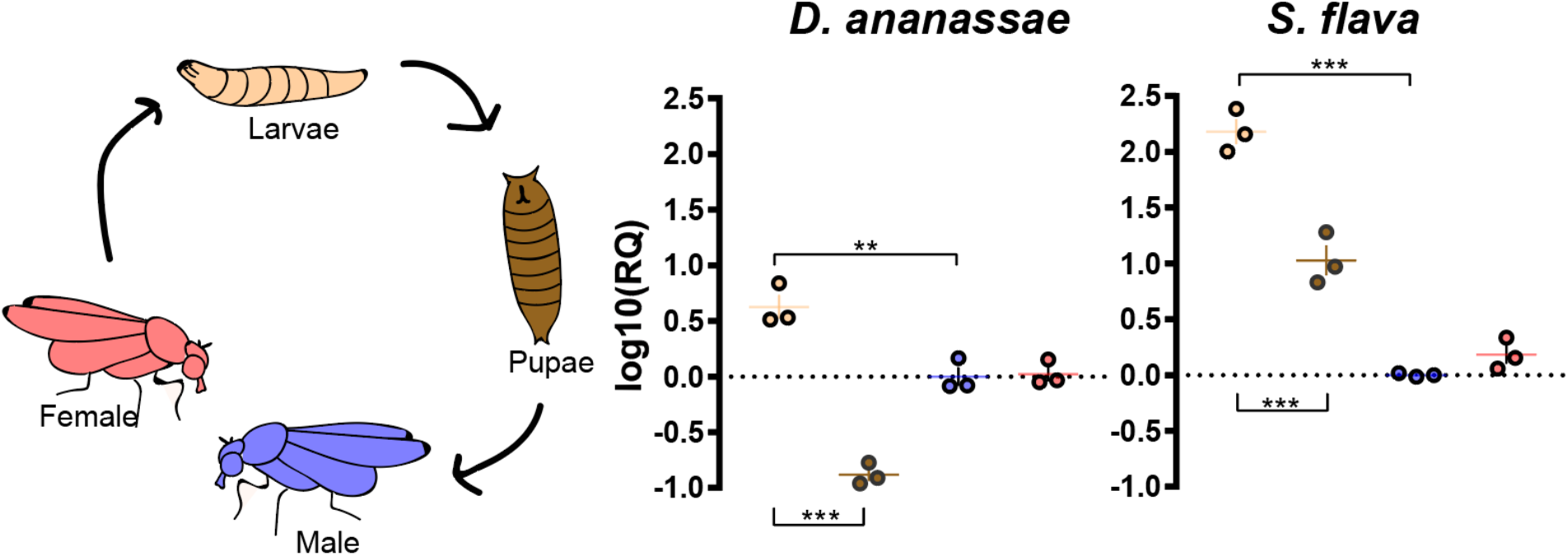
*CdtB* is expressed most highly in the *Drosophila* larval stage. Fold changes in expression of *cdtB* in two representative insect lineages (*Dr. ananassae* and *S. flava*) across development. Colors correspond to developmental stages in the left panel. Fold change is standardized against *rpl32* mRNA expression in males. Each dot represents one biological replicate. *P* <0.005**, *P* <0.0001***. All pairwise comparisons (except those between males and females) are significantly different, but are not marked for simplicity.

A critical aspect of cdtB cytotoxicity is its DNase activity, which induces double-strand breaks that can lead to cell cycle arrest, cellular distention and death (*5*). Residues in cdtB involved in enzyme catalysis, DNA binding, and metal ion binding are critical in causing mitotic arrest in eukaryotic cells and are homologous to those in DNase I. To determine if insect-encoded cdtB is a DNase, we aligned cdtB from insect lineages and other bacterial species whose DNase and cytotoxic activity are well-characterized and found that residues necessary for DNase activity are highly conserved in all insect copies (**Figure 4A, Figure S7**). To determine if these conserved residues correspond to DNase activity, we heterologously expressed and purified His-tagged cdtB from *Dr. ananassae* (**Figure S5**) in *E. coli* and utilized an agarose gel-based assay to determine its nuclease activity *in vitro* (*9*). We incubated *Dr. ananassae* cdtB (and *E. coli* cdtB as a positive control) with supercoiled plasmid pGEM-7zf(+) (Promega) at both 28°C and 37°C for 2 h. Supercoiled plasmid migrates more rapidly through a gel than nicked plasmid, which has greater surface area from relaxed superhelical tension (*4*). We predicted incubation of supercoiled (sc) plasmid with cdtB would result in a greater proportion of nicked plasmid (open coiled, or oc) isoforms. As expected, purified *Dr. ananassae* cdtB showed DNase activity *in vitro* (**Figure 4B**). Incubation at 28°C resulted in higher *Dr. ananassae* cdtB activity than *E. coli* and vice versa at 37°C, which may be a consequence of adaptation to insect and mammalian body temperatures, respectively (see **Figure S6**).

**Fig. 4.**
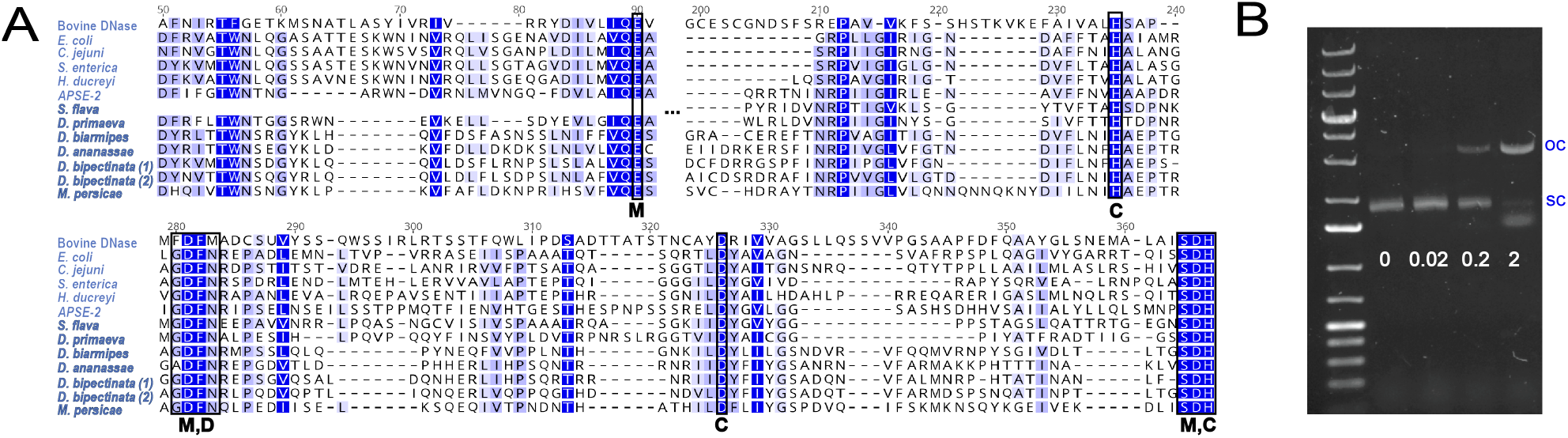
Critical DNase residues are conserved in insect-encoded cdtB and confer DNase activity *in vitro*. **A.** MUSCLE aligned amino acid sequence of DNase I and cdtB across taxa. Boxed residues are necessary for DNase activity of cdtB. Blue scale corresponds to similarity under the Blosum62 scoring matrix. Numbers correspond to alignment residue. Breaks in alignment are indicated by brackets. Species names in bold are eukaryotic. M = metal-ion binding residues, C=catalytic residues, D = DNA contact residues (*4*). **B.** Plasmid degradation following exposure to variable quantities (in µg) of cdtB from *Dr. ananassae* over 2 hrs. 0.8% agarose 1X TBE gels were stained with 0.01% SYBR(tm) Safe. OC = open-coil isoform, SC = supercoiled isoform.

The maintenance of *cdtB* in diverse insect genomes for millions of years suggests that it has an adaptive function. One clear possibility is that *cdtB* plays a role in parasitoid wasp resistance, as it does in the bacterial secondary symbionts of pea aphids (*7, 8*). Given that many drosophilid and aphid species are at high risk of parasitoid wasp attack (*27*), cdtB may facilitate protection, through DNase activity against the parasitoid wasp egg or larva. In a parasitization assay, 100% of *Dr. ananassae* and *Dr. biarmipes* survived attack by both the generalist *Leptopilina heterotoma* and specialist *L. boulardi* (*28*). It is possible, although speculative, that this unusual level of resistance is facilitated by cdtB.

To our knowledge, this is the first report of the horizontal transfer of *cdtB* from prokaryotes to eukaryotes. The domestication of *cdtB* in insects is remarkable given that the toxin originally evolved to destroy, not benefit, eukaryotic cells. Given the wealth of genetic and genomic resources available within drosophilids and aphids, horizontally transferred *cdtB* promises to be an exciting experimentally tractable system in which to explore the biology of a novel eukaryote-adapted toxin, which also has potential in targeting and killing tumor cells in humans (*29*).

## Acknowledgments

We thank Dr. Chris Jeans and Brooks Bond-Watts for their preparation of purified cdtB. Timothy O’Connor provided bioinformatics advice, Raoul O. Martin provided biochemistry advice, Julianne N. Pelaez assisted in phylogenetic reconstruction, and Anthony T. Iavarone performed mass spectrometry analysis. Coco Verster provided field assistance in acquiring specimens. Dr. Artyom Kopp and Dr. Doris Bachtrog provided *Drosophila* specimens. The *Myzus persicae* transcriptome assembly was provided by Alex Wilson and Honglin Feng. Dr. Nancy Moran ran BLAST searches of unpublished aphid genomes that corroborated our conclusions from this study. Dr. Naomi E. Pierce and Dr. Frederick M. Ausubel provided early support for obtaining a genome sequence from *S. flava.*

## Author contributions

KIV, ADG, MK, RPD, and NKW were involved in conceptualization of the project. KIV, JHW, RPD, MK, ZMA, EA, DKP and NKW conducted the investigations. KIV, RPD and NKW wrote the paper. All authors edited and approved the manuscript. **Conflict of interest:** KIV, ADG and NKW are inventors on a pending patent application related to this work, entitled “Cytolethal distending toxin B from insects for human cancer treatment”.

## Data and materials availability

*CdtB* sequences from *Scaptomyza* species and *D. primaeva* were deposited to NCBI GenBank under accession numbers MH884655-MH884659. *CdtB* codon-optimized oligos used for nuclease assays were deposited under GenBank accessions MH891796-MH891799.

## Supplementary Materials

## Materials and Methods

Materials and methods are described in the order they appear in the main text.

### Initial detection of horizontally transferred protein coding genes in insects: *Scaptomyza flava*

From the annotated genome assembly of *S. flava* (GenBank Accession RKRM00000000.1), all predicted protein sequences were queried against a local copy of the NCBI refseq protein database (downloaded May 5, 2017) using phmmer, in the HMMER3 software suite (*33*), with acceleration parameters --F1 1e-5 --F2 1e-7 --F3 1e-10. A custom perl script sorted the phmmer results based on the normalized bitscore (*nbs*), where *nbs* was calculated as the bitscore of the single best-scoring domain in the hit sequence divided by the best bitscore possible for the query sequence (i.e., the bitscore of the query aligned to itself). The top ≤ 10,000 hits were retained for further analysis, saving no more than three sequences per unique NCBI Taxonomy ID.

The alien index score (*AI*) was calculated for each query protein (modified from Gladyshev et al., 2008). The *AI* is given by the formula: AI=nbsO-nbsM, where *nbsO* is the normalized bitscore of the best hit to a non-metazoan species, *nbsM* is the normalized bitscore of the best hit to a metazoan (skipping all hits to the Drosophilini tribe NCBI:txid46877). *AI* can range from 1 to −1 and is > 0 if the gene has a better hit to a non-metazoan, which is suggestive of either HGT or contamination in the assembly. To reduce the risk of contamination, genes were considered potential HGT candidates if they were assembled on scaffolds with ≥ 5 protein coding genes and the average *AI* of the scaffold was < 0.

Phylogenetic trees of protein sequences were constructed for all potential HGT candidates with *AI* > 0. Full-length proteins corresponding to the top 200 hits (E-value < 1 × 10^−3^) to each query sequence were extracted from the local database using esl-sfetch (*33*). Protein sequences were aligned with MAFFT v7.310 using the E-INS-i strategy and the BLOSUM30 amino acid scoring matrix (*34*) and trimmed with trimAL v1.4.rev15 using its gappyout strategy (*35*). Proteins with trimmed alignments < 150 amino acids in length were excluded. The topologies of the remaining genes were inferred using maximum likelihood as implemented in IQ-TREE v1.5.4 (*36*) using an empirically determined substitution model and rapid bootstrapping (1000 replications). The phylogenies were midpoint rooted and branches with local support < 95 were collapsed using the ape and phangorn R packages (*37, 38*). Phylogenies were visualized using ITOL version 3.0 (*39*) and inspected manually to identify phylogenetically supported HGT candidate proteins. The cdtB phylogeny was the only one that passed this manual inspection.

### Identification of *cdtB* in aphid genomes and transcriptomes

An initial TBLASTN search using *S. flava* cdtB against NCBI nr resulted in hits to *Myzus persicae*, an aphid species, as well as other drosophilids (discussed below). We therefore further searched for *cdtB* in genomes and transcriptomes from representatives of Aphididae. Aphids were sampled based on availability of published or unpublished genomic resources, and included 11 species from three tribes and three subfamilies. Representatives from the subfamily Eriosomatinae, which is sister to the rest of aphids (*40*) were included in our sampling: *Pemphigus obesinymphae* and *Pemphigus populicaulis* (Subfamily: Eriosomatinae, Tribe: Pemphiginae), *Tamalia coweni* and *Tamalia inquilinus* (Subfamily: Tamaliinae), *Myzus persicae, Myzus cerasi, Diuraphis noxia* and *Acyrthosiphon pisum* (Subfamily: Aphidinae, Tribe: Macrosiphini), and *Aphis glycines, Aphis nerii,* and *Rhopalosiphum padi* (Subfamily: Aphidinae, Tribe: Aphidini). Genomes were sampled from *M. persicae* (*41*), *M. cerasi* (available on aphidbase.com), *A. pisum* (*42*), *Di. noxia* (*43*), *Ap. glycines* (available on aphidbase.com), and *R. padi* (available on aphidbase.com). We sampled published transcriptomes from the remaining aphid species (*44*–*47*).

We searched genome or transcriptome assemblies for the presence of *cdtB* with TBLASTN searches using two different cdtB proteins as the query: cdtB from *M. persicae* (XP_022163116.1) and cdtB from the *Candidatus* Hamiltonella defensa phage APSE-2 (C4K6T7), since it is infects aphid species (*48*). CdtB full or partial hits were only found in three aphids with genome sequences (*M. persicae, M. cerasi*, and *Di. noxia*), so to assess if *cdtB* was expressed in those species, we searched transcriptome assemblies for each species with TBLASTN searches using the same query proteins (**Table S1c**). For *M. persicae*, we used the assembly from the previously published transcriptome (*49*), and for *M. cerasi* and *Di. noxia* we conducted *de novo* assemblies from previously published RNAseq data. We downloaded raw RNAseq reads for *M. cerasi* (BioProject PRJEB9912, runs ERR983165 (head), ERR983166 (head), ERR983167 (head), ERR983168 (whole body), ERR983169 (whole body), ERR983170 (whole body) (PRJEB9912) and *Di. noxia* (BioProject PRJNA233413, runs SRR1999270 (whole body) and SRR1999279 (whole body) (*43*) from the Sequence Read Archive on GenBank. All runs for each species were combined into a reference transcriptome in Trinity v. 2.4.0 (*50*) using the built in Trimommatic pipeline for quality trimming (default parameters) and *in silico* normalization.

### Demonstrating *cdtB* is encoded in the nuclear genome of drosophilid species

#### *Analysis of possible contamination by coverage depth analysis in* S. flava

For *S. flava,* we aligned long PacBio reads to the genome via Burrows-Wheelers alignment (*51*) to search for unusual coverage depth relative to neighboring genes, which can be a reflection of contamination (*12*). The region containing *S. flava cdtB* did not exhibit unusual coverage depth (Grubbs’ test, p>0.05) (**Table S4**).

#### PCR and RT-PCR reaction conditions

PCR reaction conditions were composed of: 4.2 µL nuclease-free water, 7.5 µL Failsafe Premix E (Epicentre), 1.2 µL each of F and R primers (IDT), 0.8 µL DNA, and 0.12 µL of *Taq* polymerase (New England Biolabs). Thermal cycler settings were: 5 m at 95°C and 30 cycles of 95°C for 30 s, Ta for 30 s, and 68°C for 30 s, followed by 5 m of extension at 68°C. The exception to this was with *S. flava* Intergenic PCR amplification. PCR reaction conditions were composed of: 12 µL nuclease-free water, 4 µL 5X Phusion HF buffer, 0.4 µL 10 mM dNTPs, 1 µL 10 µM Intergenic F and R primers (IDT), respectively, 0.6 µL DMSO, 0.2 µL Phusion DNA polymerase (New England Biolabs), and 0.8 µL template gDNA. Thermal cycler settings were: 30 s at 98°C and 30 cycles of 98°C for 10 s, 64.1°C for 30 s, 72°C for 2 m 40 s, followed by 10 m extension at 72°C.

1% agarose 1X TBE gels were prepared with Apex Agarose in 1X TBE buffer with 1 µL SYBR™ Safe staining gel per 10 mL of gel solution. 4 µL PCR product was mixed with 1 µL ThermoScientific 6X Loading Dye and run on 1% gels in Owl ™ EasyCast ™ B1 Mini Gel Electrophoresis System rigs at 120 V for 30 m. 5 µL of ladder was used (O’Gene Ruler 100 bp or O’Gene Ruler 1kb). Gels were visualized using AlphaImager ™ Gel Imaging System (Alpha Innotech). PCR amplicons were Sanger sequenced in both directions at the UC Berkeley DNA Sequencing Facility using ABI dye terminator chemistry. Relevant gel images and primers are shown in **Figure S1** and **Table S3**.

### CdtB phylogeny reconstruction and topology test

All insect-encoded cdtB protein sequences translated from nucleotides were queried against an updated local copy of the NCBI refseq protein database (downloaded August 1, 2018) using phmmer (*33*) and default parameters, saving no more than one sequence per unique NCBI Taxonomy ID. Full-length proteins were extracted from the local database using esl-sfetch (*33*), and results from each insect cdtB search were combined to yield a final cdtB sequence set for phylogenetic analysis. Sequences were aligned with MAFFT v7.310 using the L-INS-i strategy and the BLOSUM30 amino acid scoring matrix (*34*). A total of 15 proteobacterial hits were excluded due to poor alignment and the remaining sequences were trimmed to include only the conserved cdtB domain. MAFFT was then repeated. The topology of cdtB was inferred using maximum likelihood as implemented in IQ-TREE v1.5.4 (*36*) and RAxML v8.2.9 (*52*) using empirically determined substitution models. Ten independent searches with different starting trees were carried out using each program as recommended by (*53*). The likelihood scores of all trees were re-calculated using RAxML and the tree with the highest likelihood was selected as the best cdtB phylogeny. Lastly, 1000 non-parametric bootstrap replicates were performed in IQ-TREE on the final phylogeny (**Figure S2**).

Constrained phylogenetic trees in which the insect-encoded cdtB were forced to be monophyletic were also constructed. As with the best tree, ten independent searches with different starting trees were carried out using RAxML and IQ-TREE, and the tree with the highest likelihood given the constraint was selected as the constrained cdtB tree. The best and constrained trees were then compared in CONSEL v1.2 (*54*) using the approximately unbiased (AU) test (*55*).

### Ancestral state reconstruction of *cdtB* in *Drosophila*

To construct a drosophilid species tree, DNA sequences from *adh, marf, COI, COII, 16s, cytb, gpdh, nd1*, and *nd2* (see **Table S5** for sources of phylogenetically informative genes) were aligned individually using default settings in MUSCLE (*56*) as implemented in Geneious (*57*). Alignments were visually inspected, manually trimmed and then concatenated. The final alignment included 48 species with 7479 nucleotide sites. The concatenated alignment was used to infer a drosophilid species phylogeny using maximum likelihood with the TN93 (*58*) model of nucleotide substitution in MEGA v10.0.4. The tree with the highest log likelihood (−60735.94) is shown with *Scaptodrosophila lebanonensis* as the outgroup. Branch lengths are drawn to scale and are measured in number of substitutions per site. Bootstraps values are shown (n=500) and those less than 50% were collapsed into polytomies. Maximum-likelihood ancestral state reconstruction of *cdtB* HGT occurrence was performed using the “rerooting Method” function under an equal-rates model in phytools (*59*) in R v. 1.1.456 (*60*).

### Expression of *cdtB* throughout development

#### Sample preparation and collection

Ca. 100 male and female *Dr. ananassae* (13-17 days old, the optimal egg-laying age as determined by a pilot study) were left in small embryo collection cages (Genesee Scientific, #59-100) for 6 hours with 60 x 15 mm Falcon polystyrene petri dishes filled with 3% agar in organic apple juice with a dab of Fleischmann’s active dry yeast paste. After egg-laying, eggs were cleaned and isolated in Corning Netwells inserts (#3477) and transferred onto 100 x 15 mm petri dishes with Nutri-Fly media (Genesee Scientific #66-112) prepared using standard protocols. For *S. flava,* >25 male and females (5-7 days old) were staged as above except petri dishes were filled with 3% agar and Arrowhead water with 5-9 Col-0 *Arabidopsis thaliana* leaves from adult plants submerged in the agar. For each species, we collected L2 (assessed by FBdv:00005338) (L), P-2 (assessed by (*61*)) (P), and virgin females (F) and males (M). For *Dr. ananassae* L, P, F and M, we collected 10, 5, 5, and 5 individuals, respectively, per replicate. For *S. flava,* we collected 4, 3, 3, and 3 individuals, respectively, per replicate. Samples were submerged in Ringer’s solution prior to collection. Each species and developmental stage had three replicates. Experiments occurred at 25°C under 14 h light:10 h dark cycles.

#### RNA extraction and cDNA synthesis

Samples were washed in Ringer’s solution again prior to RNA extraction. We performed RNA extraction using the Promega ReliaPrep RNA Tissue MiniPrep System following guidelines of protocol for samples < 5 mg. Final elution volume was 10 µL in nuclease-free water. RNA concentration and purity were quantified using NanoDrop ND-1000 (Thermo Fisher). We performed cDNA synthesis following standard protocols with the ProtoScript cDNA synthesis kit (NEB) using 1 µg of RNA for each sample. Synthesis of cDNA was confirmed via Qubit Fluorometer 2.0 (Invitrogen) using dsDNA HS Assay Kit (Thermo Fisher).

#### RT-qPCR primer design

RT-qPCR primers for *cdtB* and *rpl32* were designed using GenScript Real-time PCR (TaqMan) Primer Design tool (https://www.genscript.com/tools/real-time-pcr-tagman-primer-design-tool). Default primer settings were used with selection for primers. Efficiencies were determined via standard curves. Four serial 1:10 dilutions were prepared starting at 20 ng, two technical replicates and two controls with nuclease-free water in lieu of template cDNA. Melt curves showed all primer sets had high specificity. Since in most cases primers could not be designed to span exon/exon boundaries (with the exception of *S. flava rpl32*), we confirmed there was no genomic DNA contamination by loading the RNA product in a 1% agarose 1X TBE gel and by conducting RT-PCR of a no RT control and running out products on 1% agarose 1X TBE gels. Primer sequences, efficiencies, and concentrations used are shown in **Table S6.**

#### RT-qPCR cycling conditions

RT-qPCR reactions were run on StepOne ™ Real-Time PCR System (ThermoFisher Scientific). Reaction volumes were as follows: 10 µL 2X DyNAmo HS SYBR Green qPCR Kit, 0.15 µl ROX Passive Reference Dye, 0.5 µl of 40 µM forward and reverse primers, and 20 ng cDNA to a total reaction volume of 20 µl. All run cycles included initial 10 minute denaturation at 95° C, 40 cycles of: 95°C for 15 s, 60° C for 1 m, followed by a melt curve ramp from 60° C to 95° C where data was collected every +3°C. Nuclease-free water was used for no template controls.

#### Data analysis

Relative quantification was calculated according to the Pfaffl model (*62*) using primer efficiencies described in the supplement. Multiple comparisons were analyzed by two-tailed t-tests and visualized in GraphPad Prism v7.04 (GraphPad Software, San Diego, USA).

### Evaluating insect cdtB DNase activity

#### MUSCLE alignment of cdtB residues across taxa

DNase and cdtB amino acid residues were found from the following sources: Bovine DNase P00639, *E. coli* Q46669, *C. jejuni* A0A0E1ZJ81, *S. enterica* G5MJJ6, *H. ducreyi* G1UB80, *APSE-2* C4K6T7, *Dr. biarmipes* XP_016950904.1, *Dr. ananassae* XP_014760894.1, *Dr. bipectinata* (1) XP_017099970.1, *Dr. bipectinata* (2) XP_017099943.1, *M. persicae* XP_022163116.1. *S. flava* and *Dr. primaeva* sequences were translated from CDS in MH88465 and MH884659, respectively. Sequences were aligned using MUSCLE (*56*) using a maximum number of 50 iterations and visualized in Geneious (*57*) with a custom blue-scale color scheme based on the Blosum62 scoring matrix (with a threshold of 1). Thus, darker blue colors correspond to higher similarity of a residue in the alignment.

#### Cloning cdtB

*CdtB* oligos from *E. coli, Dr. ananassae, S. flava* and *Ca.* H. defensa were codon-optimized for *E. coli* expression and synthesized by GenScript Codon Optimization Services (deposited under GenBank accession #s MH891796-MH891799). *CdtB* was cloned into the pET His6 TEV vector 2B-T (a gift from Scott Gradia, Addgene plasmid #29666) using sequence and ligation-independent cloning (SLIC) (*63*).

Phobius (*64*) predicted signal peptides in cdtB from *Dr. ananassae* and *E. coli* and a transmembrane domain in *Ca.* H. defensa cdtB. In order to facilitate protein expression and purification, these domains were removed by amplifying GenScript oligos with the following SLIC-compatible primers: *E. coli* F: 5’-TACTTCCAATCCAATgcaGACCTGACCGATTTTCGTGTGG-3’; *E. coli* R: 5’-ACGACGGCTAACACCAACCGGATAGTGATCGCTGCTCATCTGGGTACGACGCGCACCATAAACAATGCCCGCTTGCAGC-3’; *Dr. ananassae* F: 5’-TACTTCCAATCCAATgcaGACGTTACCGATTACCGTATTACCAC-3’; *Dr. ananassae* R: 5’-TTATCCACTTCCAATgttattaGCCACGCGGCGCC-3’; *Ca.* H. defensa F: 5’-TACTTCCAATCCAATgcaAGCCAAAGCCACAACCACAAC-3’; *Ca.* H. defensa R: 5’-TTATCCACTTCCAATgttattaGTTAAATTTAACCGGCTTGTGGTCG-3’. For *S. flava*, SLIC was performed using the following primers: *S. flava* F: 5’-TACTTCCAATCCAATgcaATGGCGATCATTACCCGTGAGC-3’; *S. flava* R: 5’-TTATCCACTTCCAATgttattaGCCGTTCATCGGCGCC-3’. *CdtB* was cloned following University of California – Berkeley QB3 SLIC protocols (available at: http://qb3.berkeley.edu/macrolab/lic-cloning-protocol/ [accessed March 28, 2018]).

#### CdtB expression and purification

Clones were transformed into Rosetta™ 2(DE3)pLysS competent cells (Novagen) following manufacturer protocols. Freshly transformed cells were grown in 2xYT medium at 37°C to an OD_600_ of approximately 0.6, at which point the incubation temperature was lowered to 16°C. After 20 m growth at this temperature, IPTG was added to a final concentration of 0.5 mM. Cells were harvested by centrifugation after overnight growth at 16°C, resuspended in Nickel Buffer A (25 mM HEPES pH 7.5, 400 mM NaCl, 5% glycerol, 20 mM imidazole), then frozen at −80°C. Proteins were purified by Ni affinity chromatography, followed by removal of the His-tag with TEV protease, and size-exclusion chromatography.

After removal of the His-tag from the *E. coli* cdtB, a subtractive Ni affinity step was used to separate the untagged protein from the TEV protease and other contaminant proteins. The untagged protein was concentrated and loaded onto a HiPrep 16/60 Sephacryl S-200 HR size-exclusion column equilibrated in 25 mM HEPES-NaOH pH 7.5, 400 mM NaCl, 10% glycerol. Fractions containing cdtB were pooled and concentrated, assayed by UV absorption, and frozen in aliquots at −80°C.

The His-tag could not be removed from *Dr. ananassae* cdtB by TEV protease, and so the His-tagged protein was further purified by size-exclusion chromatography on a HiPrep 16/60 Sephacryl S-300 HR column equilibrated in 25 mM Tris-HCl pH 8.0, 200 mM NaCl, 10% glycerol, 2 mM EDTA, 5 mM DTT. Fractions containing cdtB were pooled and concentrated, assayed by UV absorption, and frozen in aliquots at −80°C.

We failed to purify cdtB from *S. flava* due to its aggregation into inclusion bodies. *Ca.* H. defensa cdtB was expressed at low levels and the final product contained multiple bands. Thus, these proteins were not included in the analysis of DNAse activity.

#### Mass spectrometry

Since there were several faint bands on the SDS-PAGE gel in addition to cdtB (**Fig. S5**), we analyzed trypsin-digested protein by LC-MS/MS and determined they were degradation products of cdtB and not from contaminating nucleases. Trypsin-digested *Dr. ananassae* cdtB was submitted to QB3/Chemistry Mass Spectrometry Facility at University of California - Berkeley for LC/MS analysis. 10 µg cdtB was denatured and reduced in 6.27 M urea and 9.8 mM DTT at 55°C for 20 m. Denatured protein was alkylated by incubation of 19 mM iodoacetamide for 20 m in the dark. The reaction was quenched by addition of 37.3 mM DTT and followed by overnight trypsin digestion following standard protocols for Trypsin-ultra™, Mass Spectrometry Grade (New England Biolabs). Trypsin-digested protein sample was analyzed using a Thermo-Dionex UltiMate3000 RSLCnano liquid chromatography system (LC) that was equipped with a C18 column (length: 250 mm, inner diameter: 0.075 mm, particle size: 3 µm, pore size: 100 Å) and a 1-µL sample loop. The LC was connected in-line with an LTQ-Orbitrap-XL mass spectrometer that was equipped with a nanoelectrospray ionization source and operated in the positive ion mode (Thermo Fisher Scientific, Waltham, MA). Data acquisition and analysis were performed using Xcalibur (version 2.0.7) and Proteome Discoverer (version 1.3, Thermo) software. Peptides from expressed and purified protein were measured by tandem mass spectrometry. The number of measured peptides can be used to roughly gauge the relative amounts of the different proteins in the sample. The abundance of contaminating native *E. coli* protein was negligible compared to that of heterologous *Dr. ananassae* cdtB. We searched UniProt for the contaminant, low-abundance proteins and determined none had known nuclease activity likely to lead to false positives of cdtB nuclease activity.

#### Nuclease assay

To determine DNase activity, supercoiled pGEM-7zf+ (Promega) plasmid DNA was incubated with purified cdtB. Reaction volumes were 25 mM HEPES, 5 mM MgCl_2_, 5 mM CaCl_2_ (*vis* (*65*)), 500 ng pGEM-7zf(+) incubated with variable amounts of cdtB from *E. coli* and *Dr. ananassae* in 20 µL volume. For negative controls, cdtB storage buffer was used. After 2 hours incubation in a 28°C water bath, reactions were quenched with the addition of 10 mM EDTA following protocols in (*66*). Samples were loaded onto a 0.8% agarose 1X TBE gel (premixed with SYBR Safe) and subjected to electrophoresis for 1.5 h at 90V. Images were visualized with AlphaImager ™ Gel Imaging System (Alpha Innotech).

## Supplementary Text

### Domestication of *cdtB* following horizontal gene transfer from prokaryotes to eukaryotes

The presence of eukaryotic motifs in putative HTGs after transfer from prokaryotes may indicate adaptive optimization to eukaryotic transcription, translation, and cellular function. Here we summarize how we determined if insect-encoded *cdtB* is potentially adapted to eukaryotic machinery.

#### Transcription and translation initiation elements

Thomas and Chiang 2006 (*67*) provides a comprehensive list of core promoter elements and their consensus sequences identified by transcription initiation factors (TF) TFIID and TFIIB. Other transcription or translation initiation elements we searched for included the Kozak sequence (*68*–*70*), the GC box (*71*), and the CAAT box (*72, 73*). Additionally, a Shine-Dalgarno sequence (a ribosomal binding site in bacterial RNA (*74*)), can help assess if a putative HTG is actually due to bacterial contamination.

#### Transcription termination elements

Transcription termination elements are summarized in Proudfoot 2011 (*75*) and include polyadenylation signals, cleavage site (CA), and upstream and downstream sequence elements (USE and DSE, respectively).

#### Elements of post-translational processing

Motifs involved in recognition of cargo by accessory proteins of COP and clathrin coated vesicles are described in (*76*) and were searched using the web-based database LOCATE (*77*). Additional motifs included mannosylation sites (*78, 79*), sulfation sites (*80*), nuclear localization signals (*81, 82*), and signal peptides, which were predicted using Phobius and SignalP (*64, 83*).

For the sake of brevity, we here only consider the transcriptional motifs identified bioinformatically in insect *cdtB* nucleotide sequences. This list is not exhaustive and all elements will not necessarily be found in all eukaryotic genes (*84*). We did not conduct experiments to confirm the function of the candidate elements. A visual representation of these eukaryotic motifs is shown in **Figure S4.**

Legend

- Predicted exons are highlighted light blue while predicted introns are yellow. For *Dr. biarmipes* and *Dr. bipectinata*, exons and introns were predicted by (*85*) as part of the modENCODE project. *CdtB* regions in *Dr. ananassae* and *M. persicae* were annotated based on a Gnomon gene prediction set provided by the NCBI (*D. ananassae:* FBrf0227294; *M. persicae*: LOC111028693). For *S. flava, cdtB* was annotated by aligning *S. flava* transcriptome (*86*) to an unpublished *S. flava* genome assembly.
- Coding sequence is light blue and underlined. Thus, 5’ or 3’ UTRs are light blue, non-underlined nucleotides.
- polyA signals or cleavage sites are highlighted in turquoise.
- Intergenic regions (between the two copies of *cdtB* in *Dr. bipectinata*) are lowercase.
- TATA box motifs are written in orange text, initiator sequence is in purple text, USEs are in grey text and DSEs are in blue text.

#### Dr. biarmipes

ATAAATAAGGAGAATTTCTTTCTTTTCAGTTTATTATTGAGCATCAAG<u>ATGAGAAGAATAAT</u> <u>TTTGAGCCTAGCGTTTCTGACTCGTGTAATGAGTTTAGTTACCGACTACAGACTAACGACAT</u> <u>GGAATTCTCGGGGATATAAATTACATCAAGTTTTCGATTCATTCGCTAGTAATTCATCGTTGA</u> <u>ATATTTTCTTTGTACAAGAAAGTGGAAATTTGGCCGATAAACGTTTAATTTCAATACAACAA</u> <u>AATTTACCA</u>GTAAGTTCAAGGAATATTTTTAAGTATTATACTATATTTTTTTATTATTTTCATT TTTTCTAG<u>TTTTATTTGAATGATGGTAGTAATTCTTATCTATGCGGCGCTTCTGATTTTGTGAA</u> <u>AGTGTACCAATATCAAGATCAAAGGGTTAATTTATATATATATAACTTTTTCCCGCCTCCAAA</u> <u>TG</u>GTATGACACGTTTCGTCATTATACTGATCTTTAATTAATTATATTAATAATACATTTATATT TTCTTCTGCAG<u>TTTTATCAACACTTAGTCTGACAATTGTAAGTCGAGTACCAGCCGATAGTAT</u> <u>AATTTACTTTGCTTCTCTAACGCGGCGACCTGGACGTGCATGCGAGCGAGAGTTTACAAATC</u> <u>GCCCCGTTGCCGGAATAACTATTGGCAACGATGTTTTTCTCAATATTCATGCTGAACCTACGG</u> <u>GCATCAGAAACGAAGTTCCGGATCAATTGGATGCCATTCGAAACCATATGCGCACACATGCT</u> <u>CCGCTTTCATCTTGGTTGCTAGCTGGCGATTTCAACAGAATGCCGTCATCTCTACAATTACAA</u> <u>CCATATAATGAACAATTTGTCGTCCCGCCCCTCAACACCCACGGCAACAAAATTTTGGACTA</u> <u>CCTAATTTTAGGATCTAATGACGTGAGAGTTTTTCAACAAATGGTTAGAAACCCCTATAGCG</u> <u>GAATTGTTGATCTCACTTTAACAGGTTCCGATCACAAGGCTGTACATTTTTCTCTTTG</u>AATCT CACACAATGCCACTATTTTGCCAATGTTTCAAATAATTGTAAAACTGAACACGACTAACACGATTTTTGAATAAACTCATGGTAAATTCAAA

#### Dr. ananassae

TATAGATATTTATACATGTTCCATGGCCATGTACTCATCATTCACTAAAAATTGTCCGAATCG CGCAGAACA<u>ATGAATAGAGTGCTTTCGTTATTAATTCCAGTTTTACTGAATCAGAATCTCGTT</u> <u>TCTAGTGATGTTACGGATTATAGAATAACGACTTGGAATTCAGAGGGTTATAAACTAGATAA</u> <u>AGTTTTTGACTTATTGGATAAAGACAAGTCCTTAAATTTGGTCTTAGTGCAAGAATGTGGAA</u> <u>ATATTGCAGACAAAAACCCAGGCAGCATTATTAATCCACCTGTACAG</u>GTACATAATTCATAC GTAAACAGTTTCAAGGAATAGATTTATTCAAAATGTACACTTATCTTTTTTTTCTCTAATCTG TCGAAAATAGT<u>TTATAATGATTGACGGTGAAAATGAATACGACTCTGCCAATGATGGTAATT</u> <u>ATGAAATCCGCGAGTATCGAACACGATCCACTCAATTGTTTATATATTATTTTCCGGCACCCA</u> <u>AAAGTG</u>GTAAGTATAAAATTGTTTTCAAGCGCATTTAAAATGCTGTGTGAAAAACTTGGGCA CCATGTTCTCAAGCTTACCTTGTTTGACTAGTTGTTGTAACAGTTTTTTTCTAGAAAAGATACTAAAAAGTAATTCCTAGATTCGCTTTTTTTTTAATTTTTTAATCTGTTGATGTTTCTTGACTACAAATCAATGACATAATTCAGAAACTGGAAAATTGAATCCAGGTAAAGATTAATTTCTTGGTGGAATTAATTTGCTATGAAATATTTGTTGTAATAAAATATAAAAGACAATAATGCTATATTAC ATATAAGTTTTAGTTATAAAATGTTTTATCGCACATTTTTATTTCAGT<u>TAATCAGCAATTTGG</u> <u>ATTGGCTATTGTAACCAAACAACTGGCGTCAGAGATATTATACTTTGCATCTCTTCACAATCA</u> <u>CCGAGAAATTATTGATCGCAAGGAACGTTCTTTCATTAATCGTCCTATTGTGGGATTGGTTTT</u> <u>TGGCACTAATGATATTTTTCTTAATTTTCACGCTGAACCCACTAGAAACAACGAAGTTTTACT</u> <u>TCAACTAAATGCAATTAAAACTTATATGAGCCGCTATAAACCCAATGCTTCCTGGATGCTAG</u> <u>GCGCTGATTTCAACCGCGAGCCTGGAGATGTGACTTTGGATCCACATCATGAACGATTGATT</u> <u>CACCCCTCGCAAAATACTCGCCGTAATAGAATAATAGATTACTTTATATATGGTTCTGCAAA</u> <u>TAGAAATGTTTTCGCGCGAATGGCCAAAAAACCTCATACAACAACCATAAATGCTAAGTTGG</u> <u>TCTCTGATCATAAGGCAGTAGATTTTAACCCTGCCCCGAGAGGGTAG</u>TTGGATAAGGTTGTC CAATTCCATCTTTGGCGCCCCCGGGTTAATTATAATTTAAAAAATCAAAGCAACTAGTGACAGTAACACCTTTCGAATTATAACGTATAGCCGAGTCCATGCATTTTATTTTTCGTCGTTTTTGA AAGTTTATAAAAGGCAGCCACCTTTCTTTTAATTTTGTTCAGATAAAAGCTAATCTCATTTAACATTTTGGGATTTAACTCATTTTACATGCATGCATTTTAATCTCTTATACAATTTATAATACA ATGATTTATATACAATCACTATATACAATCATTTATATACAATGAATAATAAATGATAAATGATTTTTATTTTAAATTAAATTCTTGTGTTT

#### Dr. bipectinata

AAATATCATCCCAGTACTCATCATTCACTAAACCGTCTCTGATTCGCGCGGAACA<u>ATGAACA</u> <u>CGGTGCTTTCATTAATTTTTGCGGTTCTACTGAATCGGAATCTCATTTCTGGTTTAGTTACGG</u> <u>ACTATAAAGTAATGACTTGGAACTCAGATGGTTATAAATTGCAGCAAGTTTTGGATTCATTT</u> <u>TTGAGGAACCCGTCCTTGAGTTTGGCTTTAGTGCAAGAAAGTGGAAATGTTGCTAGGCAAAA</u> <u>CCCAGGCCAAGTCATTCAACAAAATTTAGAG</u>GTACAAATACTCCTAATAAGAAAATTCAATA ATAGTATCTTAATTGCCTTTAAGTAATAGTTTCAAATACTTAATCAAAGTTTAATCCTATTTT TCATTTGAAATAG<u>TTTACTATGGCTGATGGTGAAAATGCATTTCAATGCGCCAATGATGGAT</u> <u>ATTATGAAGTCCGGCAGTATTTTAATCAAGGAACCCGTTTATATATTTATTTTTTCCCCGCAG</u> <u>CCGAAAATG</u>GTAAATACAATAATCTCTCATATTCTATCGATACGTGCTCCAATCGGATTTCCT AGTGAAGTGAATTTATACAATCACCCTTCTGAACAAATTTTTGCACGATTTTATTATACACTG ATTTAATTCAAACTTTAATTTTAG<u>TTCTACAAAAATTGGGATTGACCTTTGTAACCAGACAAC</u> <u>CGGCAACAGAGATACTATACTTCGTATCTCTGCACAATCACCAAGACTGTTTTGATCGCCGC</u> <u>GGAAGTCCTTTTATAAATCGTCCTATTCCGGGATTGGTTTTTGGCAACGACATTTTTCTCAAT</u> <u>TTCCATGCTGAACCCTCTGCAAACAACGAAGTTTTAATTCAACTACGATCCATTAAATCTTTT</u> <u>ATGAGTGTCTACAAACCCAATGCTTCCTGGATGCTAGGCGGCGATTTTAACCGCGAGCCTAG</u> <u>TGGAGTACAATCTGCTTTGGACCAAAACCATGAGCGATTAATTCACCCTTCCCAAAGGACTC</u> <u>GCCGTAATCGCATAATAGATTACTTTATATATGGATCCGCAGATCAAAATGTCTTCGCTCTA</u> <u>ATGAACAGACCTCATACAGCAACTATTAATGCTAATTTGTTCTCAGACCACTATGCTGTAGA</u> <u>TTTTAACCCTGCCCCAAGAAGAGTAGTAGGTTAA</u>TAAGTAGGTTAATAATAAATATAACTTA

CTCCACACCATACATgaattttaaactcaaacaatttataacaataaatactttgtttattaaaaattaagttctttaaattttgattttttttcaatcactaatctgaagtcgttaagtacttatcggccataagaaccctaggtcttgaaagaaatttcttcaacatttaaacttgataatagtgaaccgctcattgagttcactatcaagtgggagagtaaacaaaattctgtacaaatctggttttgattgtttgtacgagtgccgataagacaggtctgcagaaaaaaatccctcgaacaatttca ctttctggggctcgcctgaaaaatatgatcacagtactcatcattcattaaacagtTTGTGATCTGCAGAAACA<u>ATGAACACAGT</u> <u>GATTTCATTATTATTTGCCGTACTACTTAAACAGAATTTCGTTTATAGTTTAGTTACGGACTA</u> <u>CAATGTTACGACTTGGAATTCACAAGGTTATAGACTACAGCAAGTTTTGGATTTATTTTTAAG</u> <u>CGACCCGTCGTTAAATTTGGCTTTTGTGCAAGAAAGTGGAAATGTTGCTAGCGAAAACCCAG</u> <u>GAAATGTTATTCATCAGAATTTAGAG</u>GTACGGGTGGTTCGAACAAAAAACAAAAATTGGGA ATTCACCTTCTTAAGTGTATTTAATTACACATTTATTGACATAAGAGATGAAATATTATGAAA TGTGACGTGGGTACTGTGTATATTCAAGCTTTTTACTAGATCGTCGTTTTCTGAAAAGTCATT TAATTTCCAACGTAATTTTTCATATTCAATCATTTGAAATAG<u>TTTTACATAGCTGATTCTGAA</u> <u>AATGCATTTGAATGCCGCAATGATGGATATTATGAAGTACGGGAGTATGTAAATCAAGGAA</u> <u>CTCTATTGTATATATATTTTTTTCCCGCGCCTCAAAATG</u>GTGAGTACGATTATCTGTATTATTT TACCAATTTAAACTGCTGTACAATCCCAGTCTTCTATAGAAATTGTTGCCGGAATTGATTACA CTTCACATTTTTATTTCAG<u>TTCGACAAGCACTTGGATTGACTATTGTAAGCAGACATCCGGCG</u> <u>ACAGGGATACTATACTTTGTATCTCAGCATAATCACCGGGCCATATGTGATTCCCGCGACCG</u> <u>TGCATTTATTAATCGACCCGTTGTGGGATTGGTTCTTGGCACTGATGACATTTTCCTCAACAT</u> <u>CCATGCTGAACCCACTCGAACAAGGAACGAAGTACTTACTCAACTAAGAGCCGTTAGATCCC</u> <u>ATATGAGTATCCACAGACCCTCTGCTTCCTGGATGCTAGCAGGGGACTTCAATCGCTTACCT</u> <u>CAAGACGTACAACCCACTTTGATTCAAAACCAAGAACGATTGGTCCAGCCTCCACAAGGTAC</u> <u>TCACGATAACGATGTTCTGGATTACTTTATATATGGGTCCGCAGATCAGACTGTATTCGCGC</u> <u>GAATGGACAGTCCATCAAATCAAGCAACTATTAACCCAACTCTGACCGGCTCGGATCATCAT</u> <u>GCTGTTTATTTTTCCAACTATATTACATCAATTCAATACGATCGAAGCAACCTACAACATATT</u> <u>TTAGATCAAGCATGCACCGTACGAAAAACAAAGTACCACGATTGGACGAACTTTTATGATG</u> <u>AAGTTTACGATGATAAAGATAGGCTACCAAGCGCAGGTTTCATGCAGTTTCAGGACGGATCT</u> <u>TACATGCAACCCGTCGTGGTTGAACGAGATACTGATATCAAAAAGAGCTCCAATTGGTACCCGCTTGATTTGTCATTGTTAATTGGGCTTGAGGCCGCATATGCCTACAGTGTAAGGCGTAAAA</u> <u>GGAGCCTTCGTGTGTCTTCTACAACTCATCACGGCTTGATTTTCATGAACCATAAAGGCTCTC</u> <u>CAACCATGACGGTATTTTTAAGAAAAGATAAGTTGCTATGTATGACATGGACCAATAAAGG</u> <u>ATGGAGCAGTCCGACAGAATGTTATCAACACATTTCCCCTGCACAGTTTTCAGCGTCAAGGA</u> <u>ATGAACCG</u>GTATTTGACCCTTAAAATACTAAACTTTACCATTTTAATTACATTCCTTTTTCTG AATACAG<u>ACTTTTGATCGTTGTTATAACAAAGCTGAAGAATCAGTTATCAGCGAGAATTTGG</u> <u>AAACATTGGACAAAAATATCCTCAAATCCCGTAACTTGCTTATATTGCAACGTGAGTTTTCA</u> <u>AGAACCATGAGTAAGGATGGACTGTTCAACAATGAATACAATACATTTAAAGACTATTTTAA</u> <u>TGCTAAATGGAATAACACGCTAGTCAGAGATGCTTACGGCATGTCAG</u>GTAATTTACGAATAT GGATCTTAATTGTACATGTTAATTTTACTATGAATCCTTACAAAAAGTCAAACTTTTTTCTAA TTTTAG<u>GACAAGAATTCTGGGTAAAGTCCCTCAGACACTGGAATTTTGGATACTACTTCAAA</u> <u>GATGGATCTTTCTCCGTCGCATTCAGTTCTAAGCATATTCAAAACGGTGACTCCGATAATTGG</u> <u>TTGTTTATAAACGGAAATGAAATTTCCGACTGGGATCCTTGGCTCAGAAACATCTATGGAAT</u> <u>GTTTTTCTTCGACAAGTTCGGACGTCCAGTGATATTCTTTGCAGCTCTTAGTGATTACCCCCA</u> <u>TTGCTGTATCTATAGGAGCCATTGGGTGTACGAATATCGTCCAGATAATAGTTGGGATTGGA</u> <u>TCGAAAACTTTTCAGAATGGGAACCAAGCTTGTTGTCGAAAAACTCAGGCGCTTTGAGATTT</u> <u>ATTGTTGATA</u>AAAACCATATTCACAGTTCATA

#### S. flava

AGATTAACAAAGTTTTGCTGGTTTTTTTTCTATGCTTTGCCGTTGCGGCTGCTCAAAACAATTATAGATTTCTAACTTGGAACACCCAAGGACAACGCTGGCCTCAAGTAACCGCCTATTTGAATAGATACGATGTGCTATGTATTCAGGAAGCCGGTGCGTTGGGTTTGAGAGGTCTACAGGCTCTTAACCAGAATAATATAGACTATCGAATTGTGGATGAGGACAACAGAGGAGAAATTGTCACAACATCTGGTTTTAATGGTGGCGTTGAGGCTTATACGTTTTCACATGGAAACGTACCGTATTATGTCTATTATTATAACCACTTGGTGGAAACTCACGCAGCAAGGGTAGCAAACAATGACCGCA GAACGCGTAAC<u>ATGGCTATTATAACCCGTGAGCGGGCAAGCAAGGTGTACATTATTCCAGC ACATTCAAGGGATCCATATAGAATTGATGTCAATAGACCAACTATTGGAGTGAAGCTATCAGGATATACGGTCTTCACGGCCCACTCAGATCCAAACAAAAACGAAATCGTCGATACCATTGGAAAAGTGGCTCGTTTTATGGCCAGCGAAAATCAGTGCCAACAAACGAAATGGATTTTAATGGGCGATTTCAACGAAGAGCCAGCAGTTGTAAATCGACGATTGCCTCAAGCTTCAAATGGTTGTGTCATCAGTATAGTGTCTCCCGCAGCTGCCACCAGACAGGCCAGCGGCAAAATTATTGATTATGGAGTCTATGGCGGTCCACCTAGTACAGCAGGTTCGCTTCAAGCTACCACTAGAACGGGTG AGGGCAACAGTGATCATTGGCCAGTTCAAATTATGCCTGCTCCTATGAATGGCTAA</u>GAAAGC AGTTTGTTTTTGAGAAAAAAAAACATTTTTTAATAAAAAATTGAATTGAAAAAAATGTATA

Interestingly, two long terminal repeats were found flanking *cdtB* in *S. flava* (see **Supplementary Text**), which may indicate retrotransposon insertion events (*87*). Repeats in *S. flava* were identified in Geneious (*57*) function “Find Repeats” using minimum repeat length of 100 bp tolerating 10% mismatches. Two long terminal repeats were found 1,126 bp upstream of the 5’ end and 203 bp downstream of the 3’ end of *cdtB* in *S. flava;* they are 379 bp long.

Repeat 1 (5’ end):

TTTTTGGCTTGCCGTGCCGTGCTTTAATTATAATTTGGCTTCATTGCAGTTTAGCTGTTGAAAACAATACGCCAGTGGATGCTGCATTAATCATTCGAAAGCAATTTTTGCGGATTTACATGCGTTGCAGAAAACGATAATTTTCATCCATCATTTTCAAGTGTTTCCGCCAGGATTATAATACCCAGATAGACATTGCATAGCATCCTGCTGGGCTGGAACAATCCTCTTTGTGTGTTTTTCTTCTACGCTCATTATCCTTCGTTGTCGTCGTCAATTACATTGCAACATCTGAAGCTTTTAGCATTTAGTTGATGAGCTCGCCTTTTGGCGACCAACTTATCCTTCCATTCCCCGCTCCCGCCTCGAATGATTTA TG

Repeat 2 (3’ end):

TTTTTGGCTTGCCTTGCCGTGCTTTAATTATAATTTGGCTTCATTGCATTTAAGCTGTTGAAAACAATACGCCAGTGGATGCTGCATTAATCATTCGAAAGAAATTTTTGCGGATTTACATGAGTTGCAGAAAACGATAATTTTCATCCATCATTTTCAATTGTTTCCGCCAGGATTATAATACCCAGATAGACATTGCACAGCATCCTGCTGGCCTGGAACAATCCTCTTTGTGTGTGTTTCTTCTACGCTCATTATCCTTCGTTGTCGTCGTCAATTACATTGCAACATCTGAAGCTTTTAGCATTTAGTTGATGAGCTCGCCTTTTGGCGACCAACTTATCCTTCCATTCCCCGCTCCCGCCTCGAATGATTTAT G

#### M. persicae

TATATAAGGCCGACTTTTAGCGTACTGTAGGTACTATAGGTTAGGTTAGGTACATTATAATATTGCTTATTATTTATCAACATATATATATATAACGTGAATTAAAAAAAATTTAACATTTAAAA CTATAAGTTAGAAGCTGAAATAATATCTACAGA<u>ATGGCGACAATAGTCTTGCTATTATTAAT TTCTCAGCTTATAAATTATAATTTAATTTCGTGTTTAGTTACTGATCACCAAATAGTAACTTGGAATTCAAATGGCTACAAACTCCCAAAAGTTTTTGCTTTCTTGGATAAGAATCCACGTATACATTCAGTTTTTGTGCAAGAAAGTGGAAATGTTGAATCTGAATCAAATAATGCAGGAACTCCA GTACCGAAAACTAATTTACCAAAA</u>GTATAAGGATTTTTAATTTTGATACAATTATTGTATAAT ATATTTAAATATTTGATATTTTTTAATTTTTTAAAATATTTTAATTTATAGTTTAATGTCTCAA ATATAATTTTTTTATATTTAATTAAATATAATAATCATATTTTTATAATTATTTTATATTAAAA AAAATGATTTTTTTAAATAG<u>TTTGTTATTGCTGACGTAGAAGGTGATATCGAATGTACGAATT ATGCTGACTTTATTAAAGTCAAAACTGTAACGACATTTCAAAACGGGCCGTTTTATATATATT ATTCACCTGCATCCCCCAATG</u>GTAAGTAACATCAAATAATTTCAATAATACCTACAATTTTCA TATTATTTTGTACTTCTATTATCTAG<u>TTACTCCCAAAATTAGCATAACTATTGTAAGCAGACA TTTAGCTAGAGAGATAATACTTTTTCCAAGTCAACACAATCATAAGAGTGTATGTCACGATCGTGCTTATACTAACCGCCCTATTATTGGATTGGTTTTACAAAATAATCAAAATAATCAAAAAAATTATGATATTATTTTAAATATTCACGCGGAACCAACTAGAAAACGTAACGAAGTGATAACACAATTGAAAATTATCAGAACTTATATGAATACCATTAGAAAACCTACTTCATGGTTGTTAGCCGGTGATTTTAATCAATTACCTGAAGACATTATAAGTGAATTAAAATCTCAAGAACAAATAGTCACACCAAATGATAATACTCATGCTACTCATATTCTCGATTTTTTAATATATGGTTCCCCTGATGTTCAAATTTTTTCGAAGATGAAAAATTCTCAATATAAAGGAGAAATAGTGGAAAAGG ATTTGATTTCAGATCATAAAGCCGTGCATTTTTTTAAGTAG</u>TTAGGTAATTTTTTATTAATGTT TATTATTTTGGTGTTAGCGCATTGTATTCATCAAATATTTGTGAATTTTATTATTAAATTTATTTTTATTTAACATAATAATTTATTAAGTAGGTACCTATATTTTACTATGGTCTGAATGACTGAATTTTGATGATTGATGATTGTCTATAATATAATATGATAGCCATTACACCATATTCAAAATAATAAGTACCTATTGTATATTTAAATAATTAAATTATATTTCATAGGTAACATGTATTTTTTAAGTTATTTATTTAAGAATTTTAAATGTATTTTCACTCACAAAAAAAACAAATATGAATTGTACAATTGAGAAAACTCGGGGTCATAGAATAATCATTTTTGATAAAATAAATCGTTTATAATATTTTACTGTATACCGACTATTATAATTTGATAATTAATGTATTGTC

### Apoptosis inducing protein 56 is fused at the C terminus with cytolethal distending toxin B in *Dr. bipectinata* and other ananassae subgroup species

*Apoptosis inducing protein 56 is fused at the C terminus with cytolethal distending toxin B in* Dr. bipectinata *and other ananassae subgroup species.* The full cdtB alignment (**Figure S7**) shows a large C-terminal region of the second *Dr. bipectinata* copy (with five exons) does not align to that of any other cdtB sequences except two other species in the ananassae species subgroup, *Dr. pseudoananassae nigrens* and *Dr. ercepeae.* To determine if this region was a bacterial contamination artifact, we amplified and sequenced this region from *Dr. bipectinata* via RT-PCR and Sanger sequencing, which confirmed that this region had introns (leading us to disfavor bacterial contamination) and overlaps the *cdtB* domain (disfavoring errant colocalization of the two genes by assembly error), which corroborated data from (*85*). We extracted this conspicuous region (residues 294-651 from XP_017099943.1) and submitted it as a BLASTP query **(Table S8a),** which showed the region has high homology to another *Ca.* H. defensa protein, hypothetical protein D (*88*). Interestingly in the *Ca.* H. defensa 5AT strain genome (NC_012751.1) the two genes *cdtB* (KF551594.1) and *ORF D* (DQ09613.1:2421-3185) are ca. 255 bp apart. Since we identified relatively few hits in the first BLASTP search and high divergence may have limited the number of identified homologs, we subsequently ran a BLASTP search using the *Ca.* H. defensa hypothetical protein D (WP_015874047.1) as a query (**Table S8b**). Our results show that the second half o*f D. bipectinata* cdtB has homology to the protein apoptosis inducing protein 56 (aip56), a key virulence factor of *Photobacterium damselae piscida*, one of the most important bacterial diseases in mariculture (*89*). Aip56 is secreted by the type II secretion system of *P. damselae piscida* (*90*). In infected fish, aip56 triggers apoptosis of macrophages and neutrophils, which leads to infection-associated necrotic lesion that can devastate a population (*91*).

#### Aip56 may have been transferred horizontally in other eukaryotic species

Interestingly, copies of *aip56* were also found in the genomes of two other insect species: *Operophtera brumata,* the winter moth, and *Danaus plexippus plexippus,* the monarch butterfly. However, the two *aip56* homologs (OWR44524.1, OWR45007.1) identified in *Da. plexippus* were located on short scaffolds containing only the gene of interest, suggesting these may be bacterial contaminants. While *aip56* homologs found in *O. brumata* were also found on relatively short scaffolds (1.6kb – KOB51764.1; 52 kb – KOB69574.1; 48kb – KOB68847.1, KOB68849.1) the two former identified scaffolds encoded other *bona fide* insect genes. Additionally, the latter scaffold had two *aip56* orthologs arranged in tandem, one of which contained an intron. Here, we only considered aip56 homologs to be insect-encoded if they were found on scaffolds containing other *bona fide* insect genes. Thus, we include *Dr. bipectinata aip56* and at least three *O. brumata aip56* copies. A previous study determined via phylogenetic network analysis that eukaryotic aip56 sequences cluster together, and are closely related to APSE-2 hypothetical protein D (*92*).

#### Aip56 is an AB-toxin, but only residues in the B domain appear to be conserved between eukaryotic and prokaryotic species

Aip56 is an AB-toxin (*89*). While relatively little is known about aip56, motifs have been discovered that facilitate their cytotoxicity, which manifests as induction of apoptosis in eukaryotic cells (*93, 94*). Aip56 is composed of two domains linked by a disulphide bridge: the A domain is responsible for catalytic activity and the B domain facilitates cellular entry of the toxin. One of the key components of the A domain is a HEXXH motif which is typical of zinc metalloproteases and is highly conserved within bacterial species (*95*). The A domain cleaves the transcription factor NF-KB 65, thus interfering with the regulation of inflammatory, anti-apoptotic genes (*95*), affecting bacterial pathogenicity. In an alignment of all aip56 sequences we identified in **Table S8,** an HEXXH motif was found in all bacterial species except in *Ca.* H. defensa, and was absent in all insect-encoded copies (**Figure S8**). In the B domain, unlike the A domain, we could identify several motifs that appeared to be conserved between eukaryotes and prokaryotes (e.g. FD^695-696^, GRP^698-700^). While the mechanisms behind B domain cellular entry are less defined, it is known that a deletion of this delivery module domain inhibits binding to target cells and reduces cytotoxicity, making it plausible that these residues are important in cellular uptake (*95*). Given that this domain is vital in cellular entry in diverse hosts (*89, 91*), their conservation across domains of life may signal a vital role in facilitating clathrin-dependent endocytosis, the mechanism of aip56 and cdtB uptake (*91, 96*). Both cdtB and aip56 also undergo endosomal maturation in host cells prior to inducing cytotoxic effects (*91, 97*), suggesting compatibilities in their methods of cytoplasmic delivery that may have facilitated the *cdtB+aip56* fusion in the ancestor to *Dr.* ananassae subgroup species.

#### Hypotheses on the history, functions and mechanisms of cdtB + aip56 *fusion protein*

The fusion of *cdtB* and *aip56* in *Dr. bipectinata,* along with the proximity of those two genes in the *Ca.* H. defensa genome, strongly suggests that these two genes may have been horizontally transferred between ancestors of *Ca.* H. defensa APSE-2 and eukaryotes, either directly from a phage or via a bacterial intermediate (*98*–*100*). These two genes, which are encoded in an operon-like fashion in *Ca*. H. defensa, are close enough (within 300 bp) that small mutations could have led to read-through mutations or frameshift mutations and to the two individual proteins being expressed and translated as one larger protein (*101*). CDTs and aip56 are both AB toxins, and in *Dr. bipectinata* + *Dr. pseudoananassae* only the A subunit of the CDT, encoded by *cdtB*, and the B subunit encoded by *aip56* are found in these species. Thus, it is plausible that this ctdB+aip56 fusion was adaptive in some insect backgrounds because these two protein domains can work in concert to affect cellular internalization (aip56 B domain) followed by DNase and apoptogenic activity (cdtB). We speculate that the shared cellular internalization pathways of both cdtB and aip56 (clathrin-mediated endocytosis and endosomal maturation) are in fact synergistic. We hypothesize that the *cdtB+aip56* fusion represents a unique adaptation to the same problem *cdtB* may have evolved in response to. However, it is also clearly not the only viable way of affecting cdtB function, since we could not find *aip56* in *Dr. ananassae, Myzus* spp., or *Dr. primaeva* + *Scaptomyza* spp. genomes.

### Analysis of *cdtB* synteny within drosophilid and aphid lineages

In order to assess the evolutionary history of horizontal transfer in *cdtB* within and among insect species, we analyzed synteny of *cdtB* in aphid and drosophilid species (*26, 102*–*105*). We downloaded *Scaptomyza, Drosophila* and Macrosiphini aphid scaffolds and compared gene identity up- and down-stream of *cdtb* in SynMap using CoGe (*106*). SynMap revealed clear *cdtB* synteny between species in each of these three clades: *Dr. bipectinata* + *Dr. ananassae, Dr. primaeva* + *S. flava, Di. noxia* + *M. persicae* + *M. cerasi,* and none between *Dr. biarmipes* and any other species.

Additionally, since variability in scaffold size between species could limit syntenic inference, we used a complementary microsyntenic approach and manually identified genes flanking *cdtB* (see **Table S9**) in representative drosophilids, which corroborated the results from CoGe.

Comparison of *cdtB*-containing scaffolds to those in *Dr. melanogaster* indicate that *cdtB* is located on Muller element E (chromosome 4) in *S. flava* and *Dr. primaeva*. In contrast, *cdtB* is located on Muller element B (chromosome 3R) in *Dr. ananassae* and *Dr. bipectinata* as well as in the more distantly related *Dr. biarmipes* (*107*).

Results for the above SynPlot analyses can be regenerated at the following links:

- *Dr. ananassae* to *Dr. bipectinata*: https://genomevolution.org/r/qc7l
- *Dr. ananassae* to *S. flava*: https://genomevolution.org/r/s735
- *Dr. ananassae* to *Dr. biarmipes:* https://genomevolution.org/r/s3ze
- *Dr. bipectinata* to *Dr. biarmipes*: https://genomevolution.org/r/s3zk
- *Dr. biarmipes* to *S. flava*: https://genomevolution.org/r/s737
- *S. flava* to *Dr. primaeva*: https://genomevolution.org/r/135zl
- *M. persicae* to *M. cerasi*: https://genomevolution.org/r/12l4j
- *Di. noxia* to *M. persicae*: https://genomevolution.org/r/139ic

**Fig. S1.**
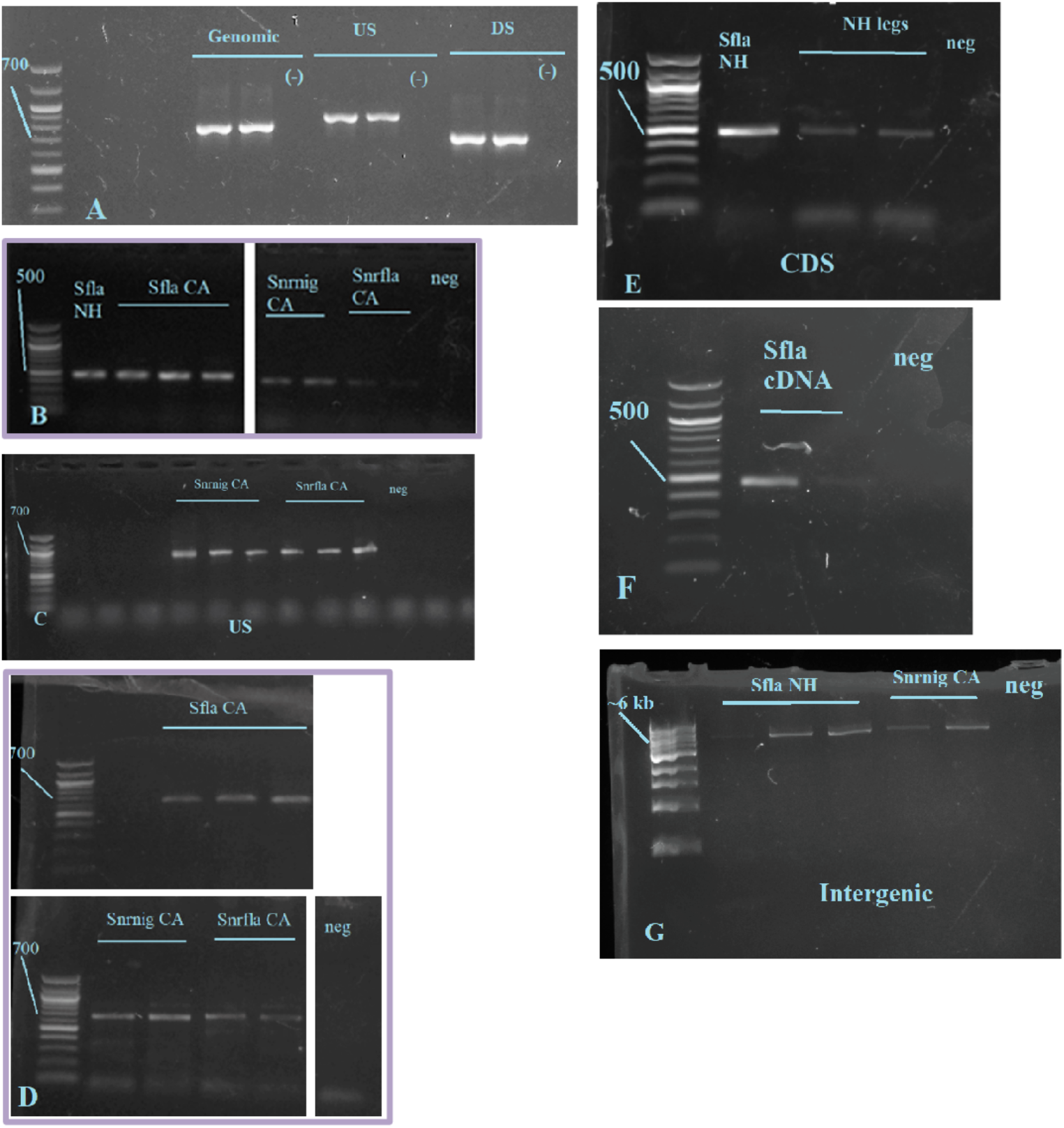
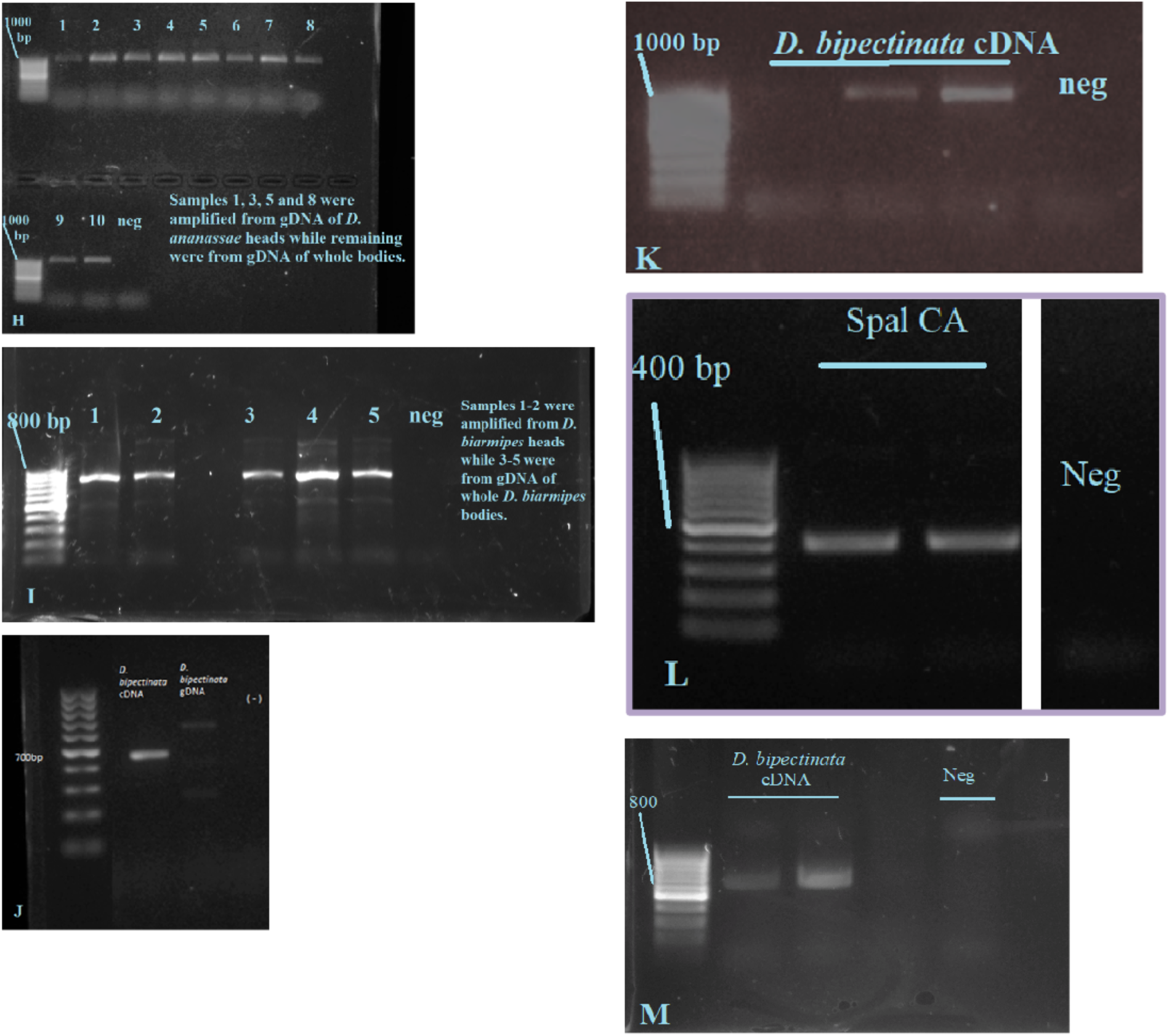
Gel images of amplicons using primers from **Table S3.** Throughout, *Sfla* NH refers to *S. flava* colonies originally captured from New Hampshire, USA and maintained at UC-Berkeley. *Sfla* CA refers to *S. flava* collected from Berkeley, CA that are 100% identical at *COI* to *S. flava* from NH. *S*nr*nig* CA refers to *S.* nr. *nigrita* captured from Berkeley, CA. *S*nr*fla* CA refers to *S.* nr. *flava* collected from Berkeley, CA. *Spal* refers to *S. pallida.* ‘Neg’ or (-) refers to negative controls with nuclease-free water substituted for template DNA. Unless otherwise specified, *cdtB* was amplified from single whole bodies of the drosophilid species. In all figures O’Gene Ruler 100 bp Ladder (ThermoFisher) is used in the first well (except for item G, in which O’Gene Ruler 1kb Ladder (ThermoFisher) is used). Numbers adjacent to the ladder are approximate size in bp. 1% agarose 1X TBE gels were stained with 0.01% SYBR™ Safe. Amplification from heads and/or legs in E, H and I indicates that *cdtB* is unlikely to be from contamination of gut bacteria. In images B, D and L, different images from the same gel and primer set are stitched together for clarity, these divisions are indicated by white lines. Besides these concatenations, these images are not subject to any nonlinear adjustments, and are indicated by a purple border.

**Fig. S2.**
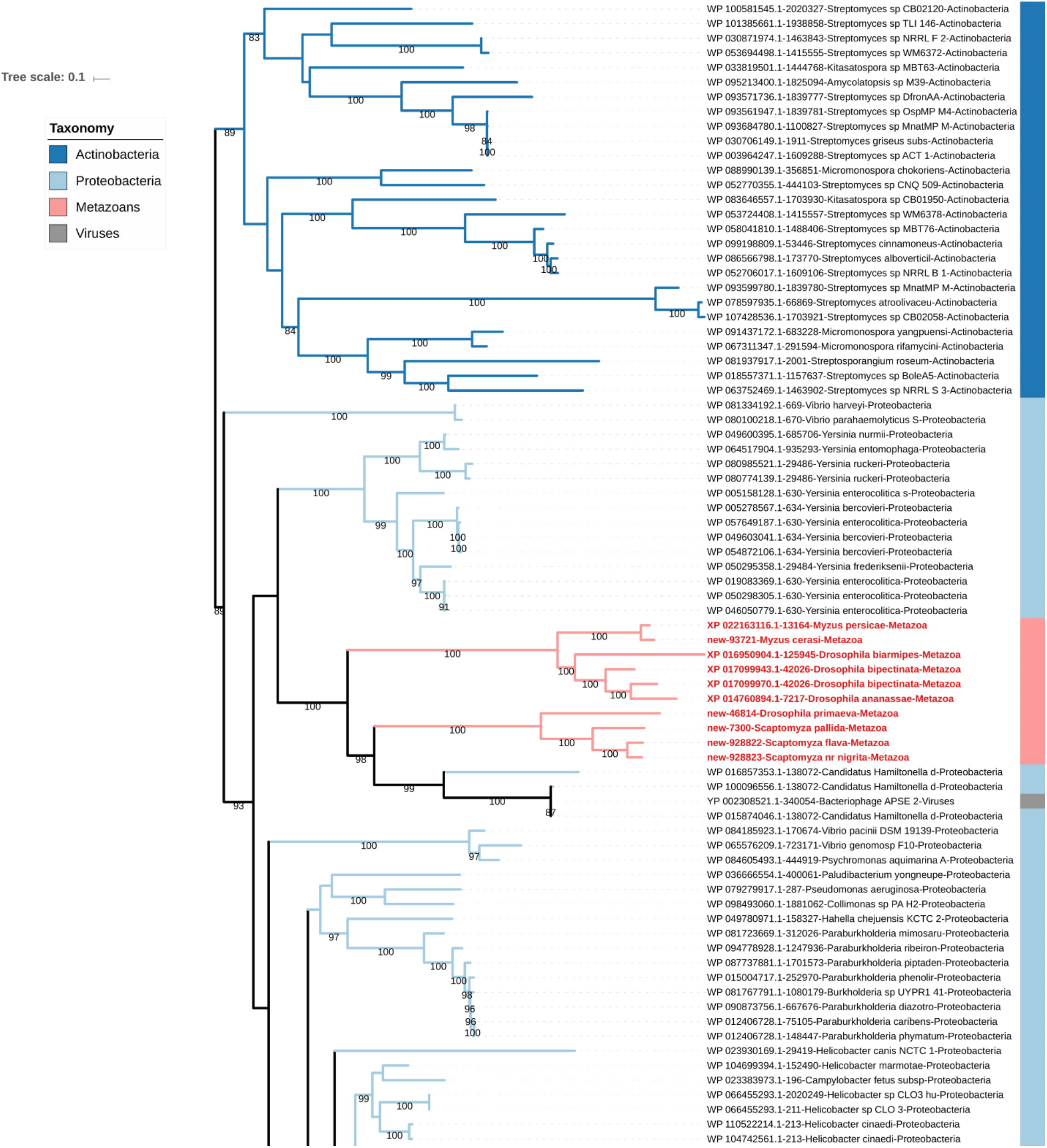
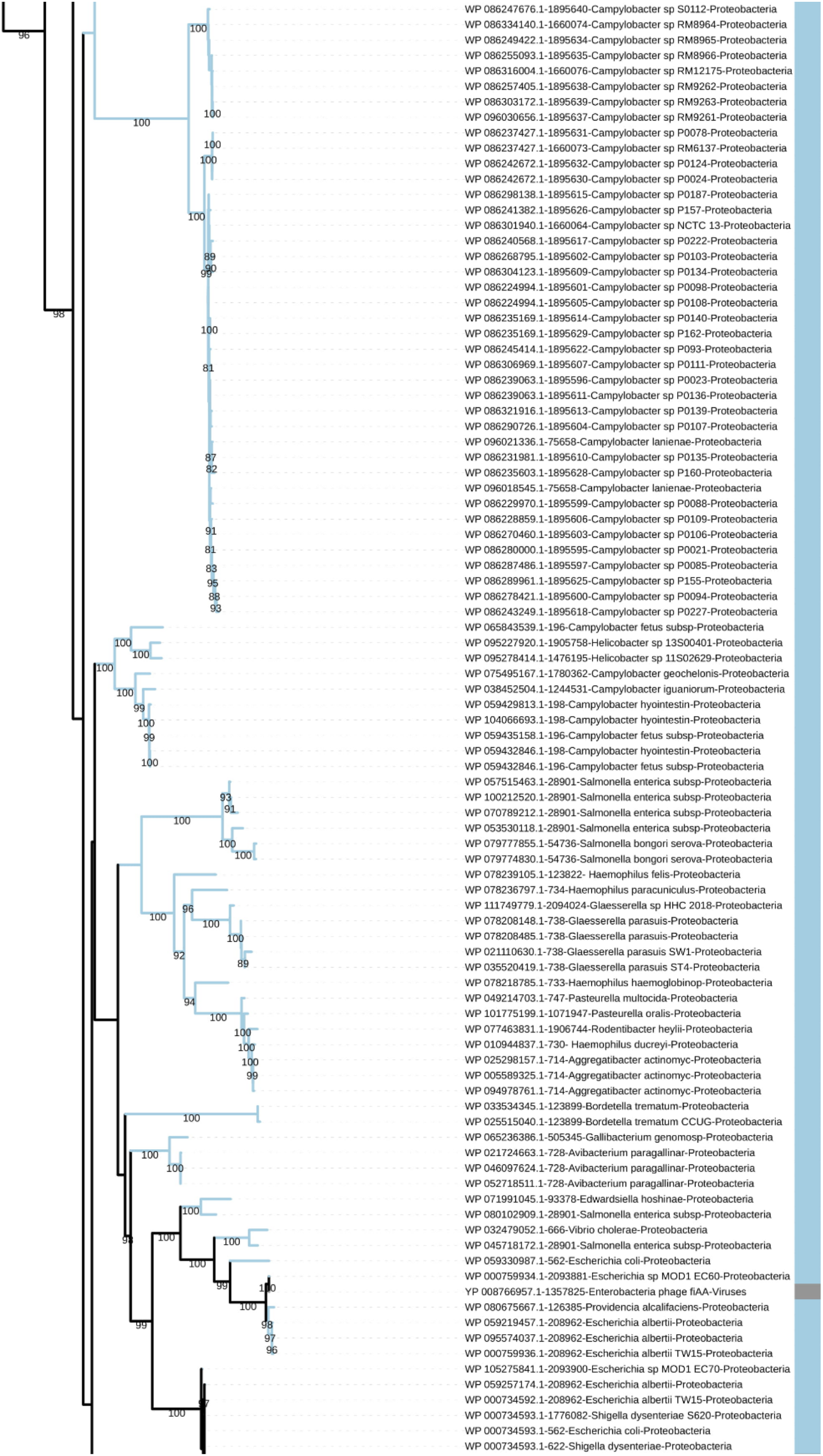
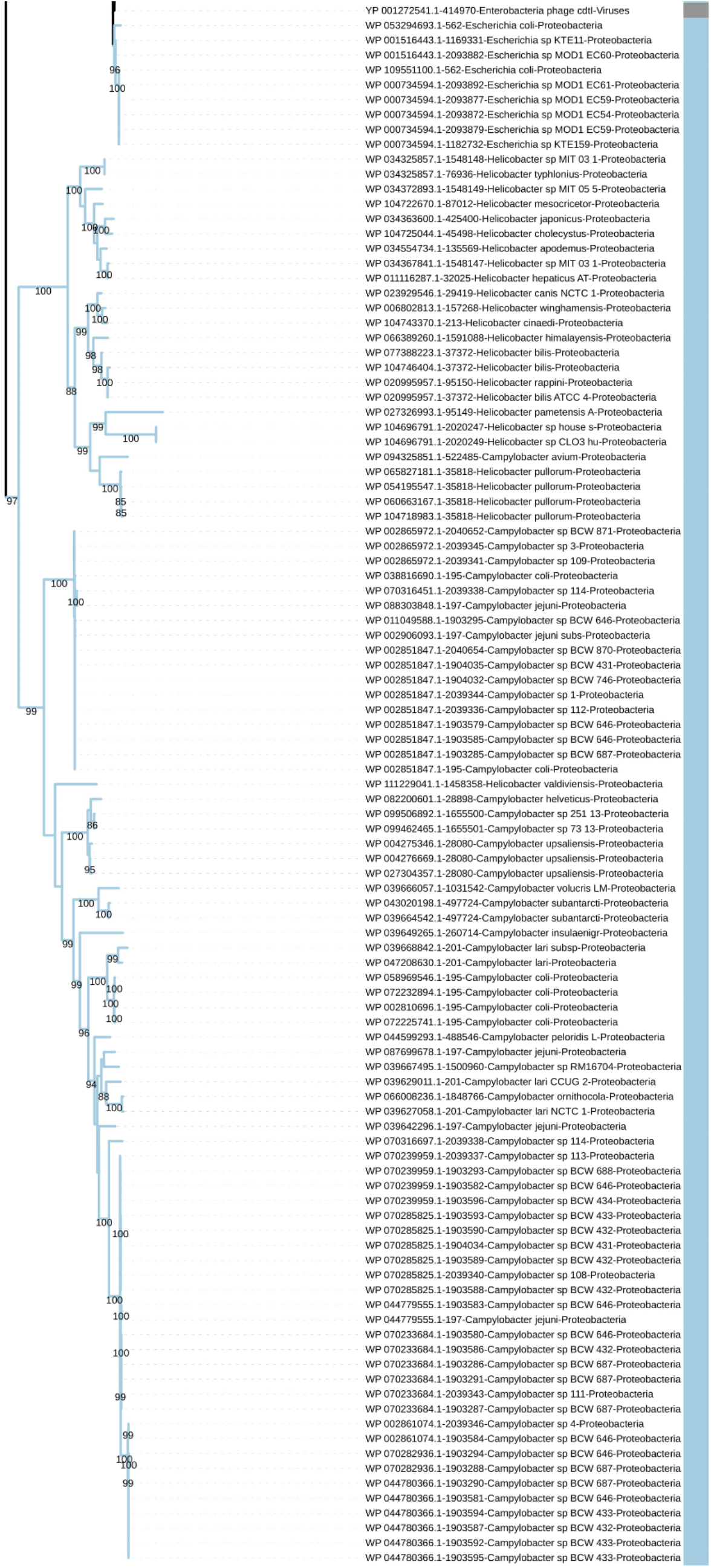
CdtB protein maximum likelihood phylogeny incorporating sequences from NCBI refseq protein database and those found in this study. Numbers on branches are support values from 1000 bootstrap replicates. The names at the tips indicate the NCBI Protein ID (if not found in this study, in which case it is described as ‘new’), species name and domain. For a further description of this phylogeny, please see **Materials and Methods**.

**Fig. S3.**
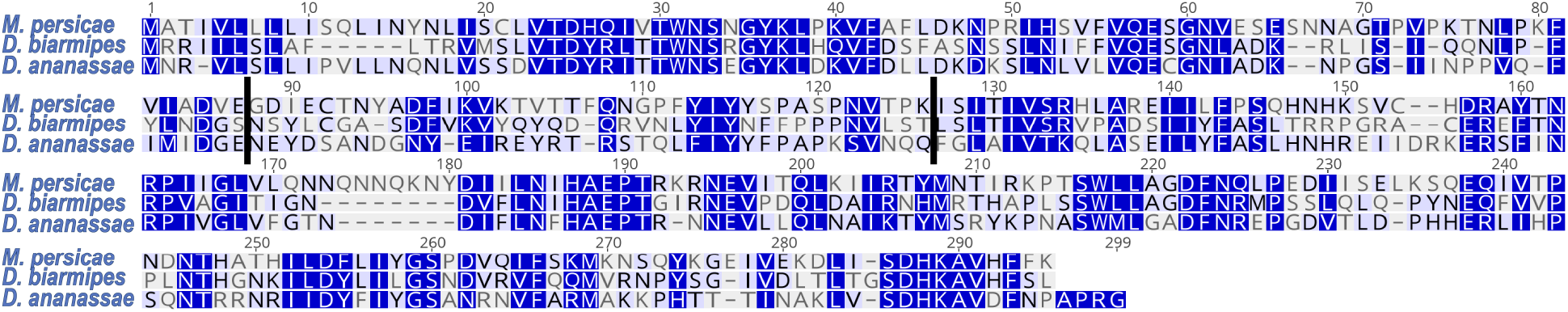
MUSCLE cdtB amino acid alignment for representative intron-containing copies of cdtB. Splice junctions (indicated by black lines) are conserved in a MUSCLE alignment cdtB copies from *Dr. ananassae, Dr. biarmipes* and *M. persicae.* Blue scale corresponds to similarity under the Blosum62 scoring matrix with a threshold of 1 (where darker shading corresponds to higher similarity). Numbers indicate alignment residues.

**Fig. S4.**
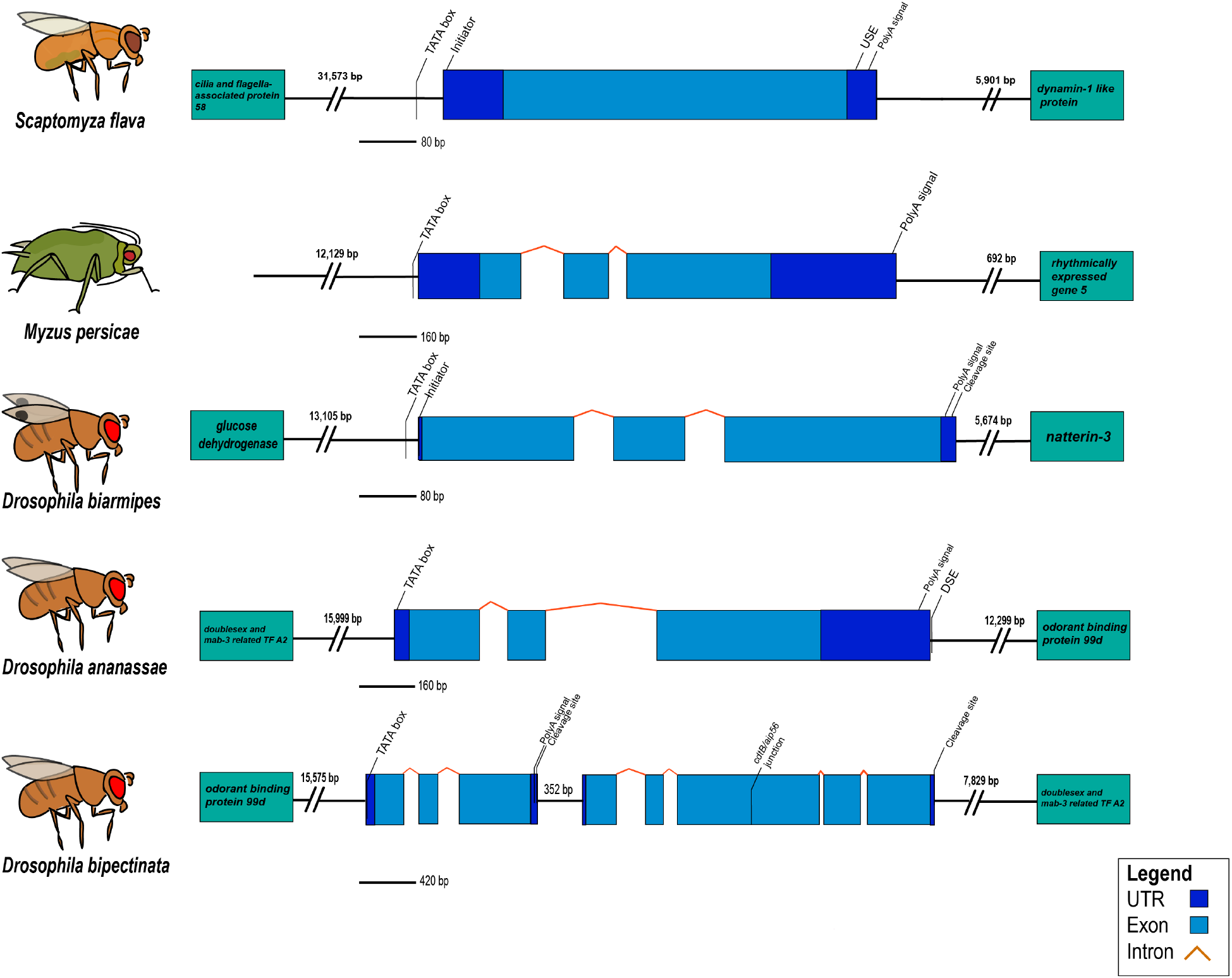
Gene region and eukaryotic motifs of *cdtB* in representative insect species. UTRs are 5’-3’ left to right. Dark blue boxes are UTRs, orange bent lines are introns, and light blue boxes are exons. Boxes to the left and right are nearest flanking genes and brackets indicate distance to nearest gene. Slanted lines with floating text indicate motifs described in the **Supplementary Text**. Gene representations are drawn approximately to scale with calibration legends.

**Fig. S5.**
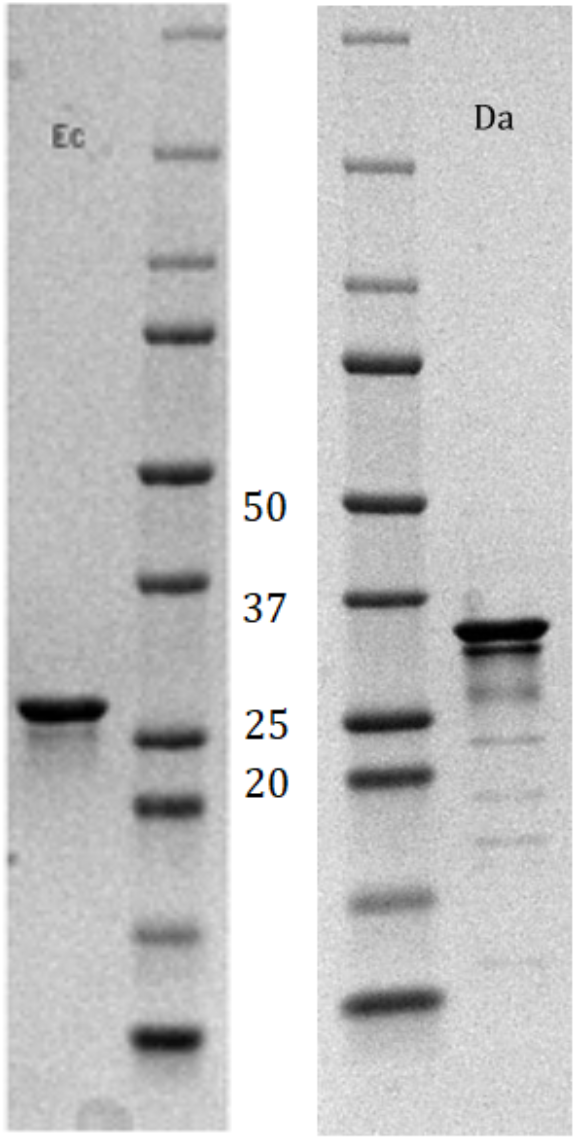
CdtB from *E. coli* (Ec) and *Dr. ananassae* (Da) washed, separated by 10% SDS-PAGE, and visualized by staining with Coomassie blue. Molecular mass markers are in kDa.

**Fig S6.**
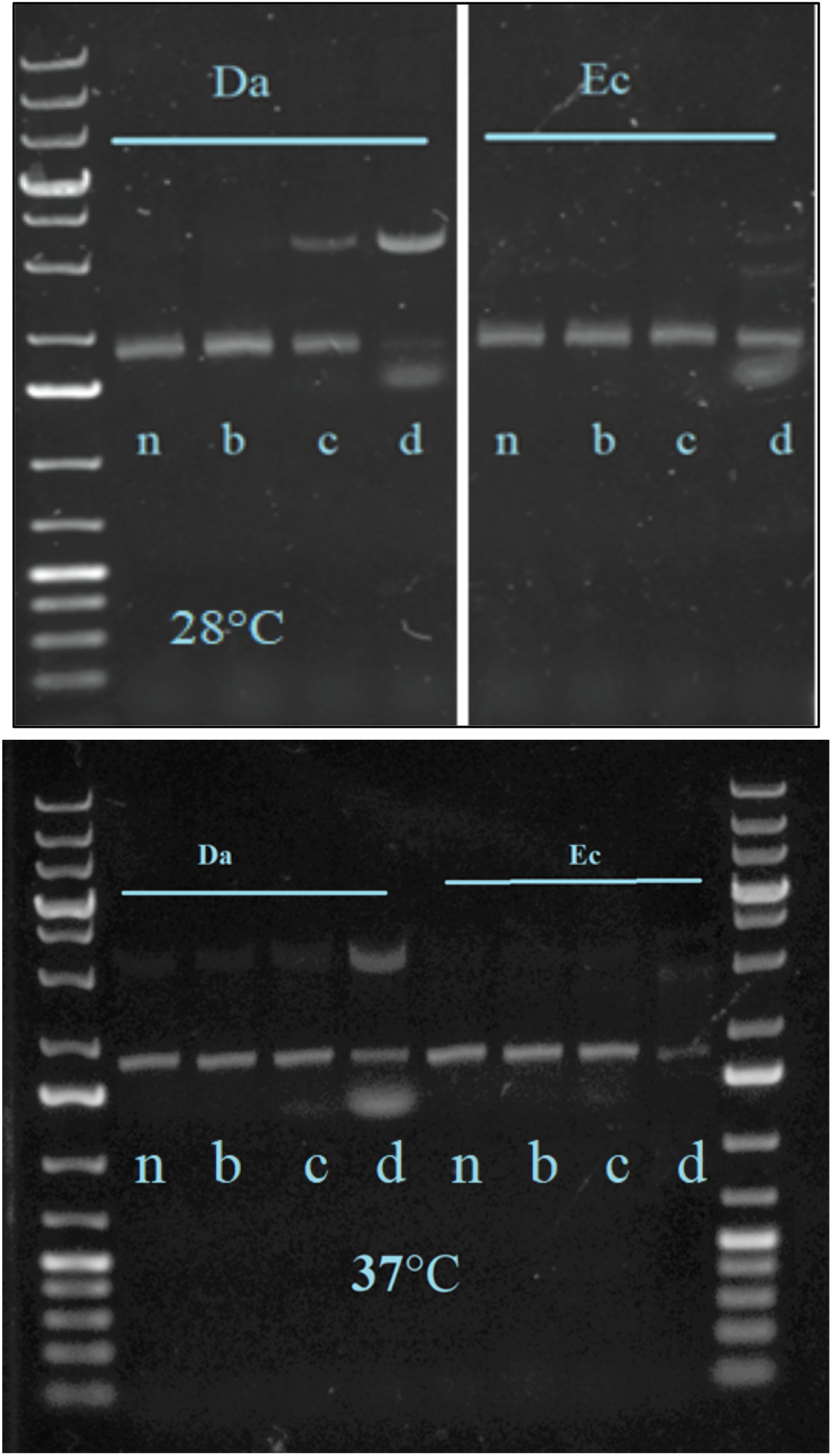
DNase activity assays of *Dr. ananassae* (Da) cdtB and *E. coli* (Ec) cdtB for 2 hours at two different temperatures, 28°C and 37°C. n = buffer control (no cdtB), b = 0.02 µg cdtB, c = 0.2 µg cdtB, d = 2 µg cdtB. 0.8% agarose 1X TBE gels were stained with 0.01% SYBR™ Safe. 5 µL of O’Gene Ruler 1kb Ladder (ThermoFisher) are in the first and last wells in the image. For clarity, the first image is stitched from two parts of the same gel, and this division is indicated by a white vertical line. There were otherwise no vertical manipulations or nonlinear adjustments. For information on incubation conditions refer to the **Materials and Methods.**

**Fig. S7.**
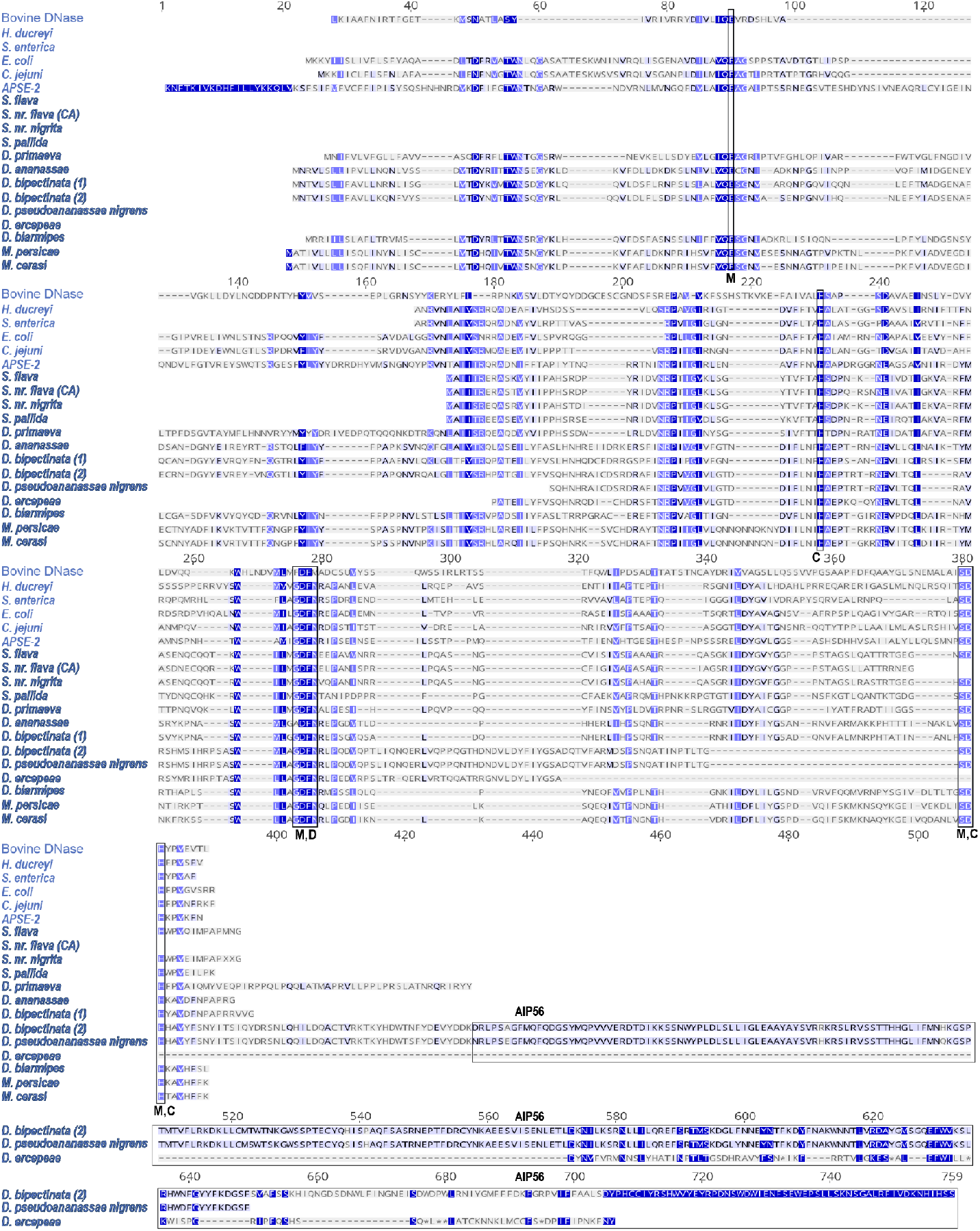
MUSCLE alignment of cdtB from all identified eukaryotic species with non-pseudogenized *cdtB* copies. Vital DNase residues are highlighted in blue. Blue scale is based on BLOSUM62 similarity scores where darker residues are more similar. Bold species are eukaryotic. DNase and cdtB amino acid residues were from the following sources: Bovine DNase P00639, *E. coli* Q46669, *C. jejuni* A0A0E1ZJ81, *S. enterica* G5MJJ6, *H. ducreyi* G1UB80, *APSE-2* C4K6T7, *Dr. biarmipes* XP_016950904.1, *Dr. ananassae* XP_014760894.1, *Dr. bipectinata* (1) XP_017099970.1, *Dr. bipectinata* (2) XP_017099943.1, *M. persicae* XP_022163116.1. *Scaptomyza* spp. and *Dr. primaeva* were translated from CDS in GenBank sequences MH884655-MH884659. *Dr. pseudoananassae nigrens* and *Dr. ercepeae* sequences were translated from sequences found from their transcriptomes (*11*). *M. cerasi* sequence acquisition is detailed in **Methods.** Residues vital for DNase activity are described in the main text. AIP56 domain is indicated in a black box.

**Fig. S8.**
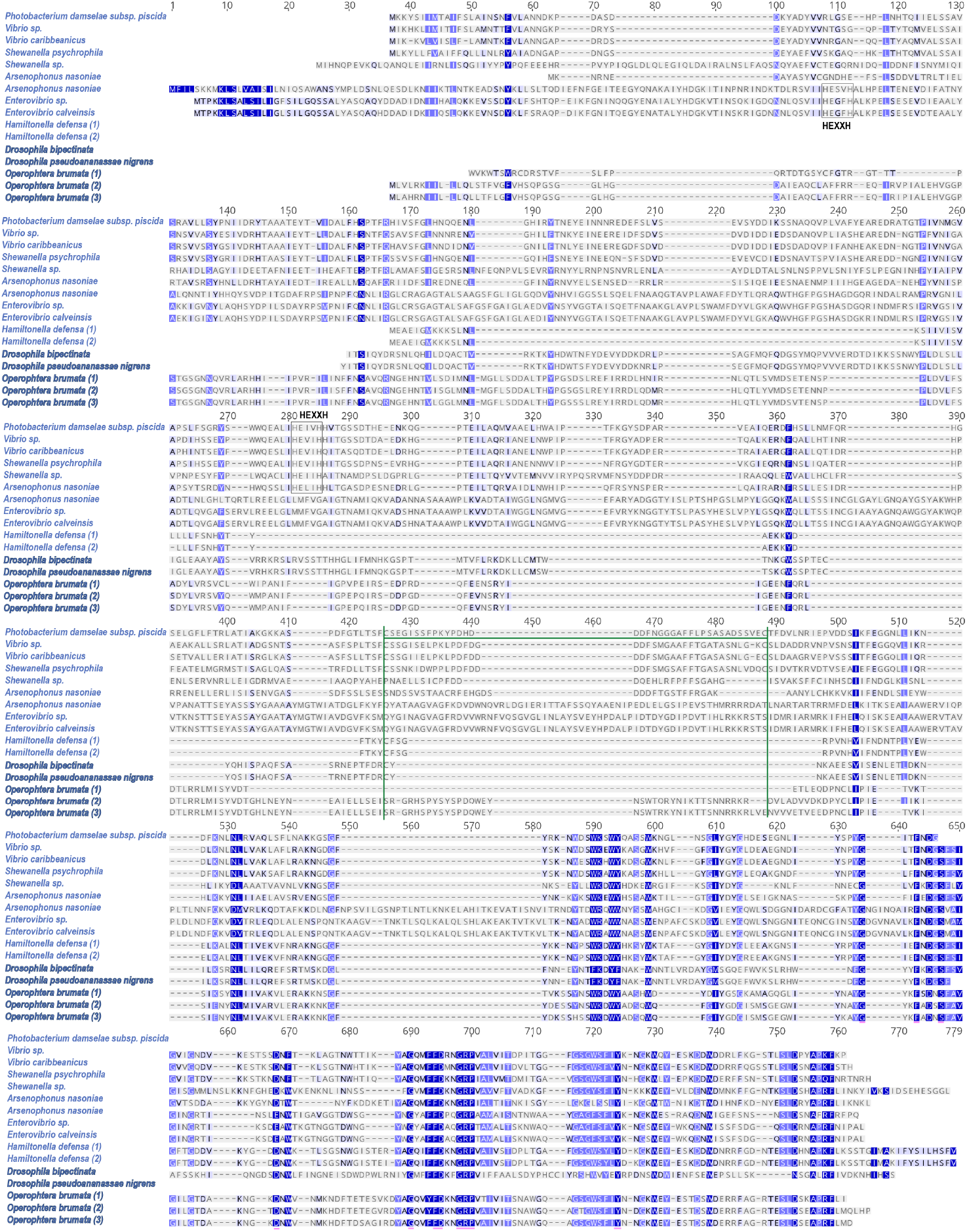
MUSCLE alignment of aip56 from representative bacterial species and insect species. Bolded species re eukaryotic. Green borders indicate a disulfide bridge (green line) that separates the A (N-terminal) and B (C-terminal) domains. The HEXXH motif in bacterial species except *H. defensa* are boxed and labelled. Sequences were found from the following sources: *P. damselae* subsp. *piscida*: WP_094461508; *Vibrio* sp.: WP_089070319; *Vibrio caribbeanicus:* WP_009600485; *Shewanella psychrophila:* WP_077754668; *Shewanella* sp.: WP_012326868; *Arsenophonus nasoniae*: WP_051297127, WP_051296919; *Enterovibrio* sp.: WP_102315974; *Enterovibrio calviensis*: WP_017014894; *H. defensa* (1, 2): WP_015874047, WP_100096555, respectively; *Dr. bipectinata* ‘tail’: XP_017099943.1 residues 294-651; *Dr. pseudoananassae nigrens*: extracted and translated from (*11*); *O. brumata* (1, 2, 3): KOB68849, KOB69574, KOB68847, respectively.

**Table S1.**
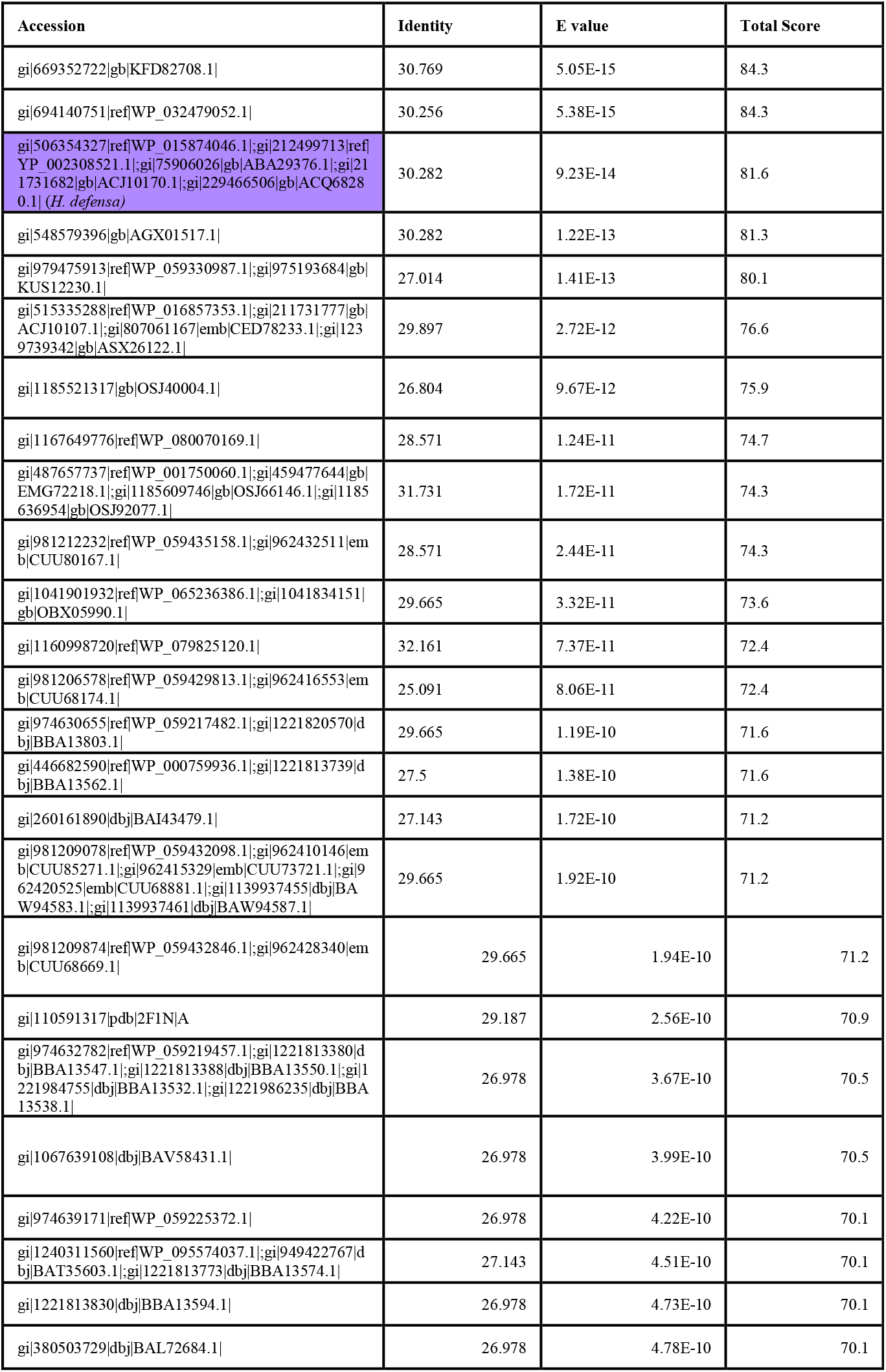
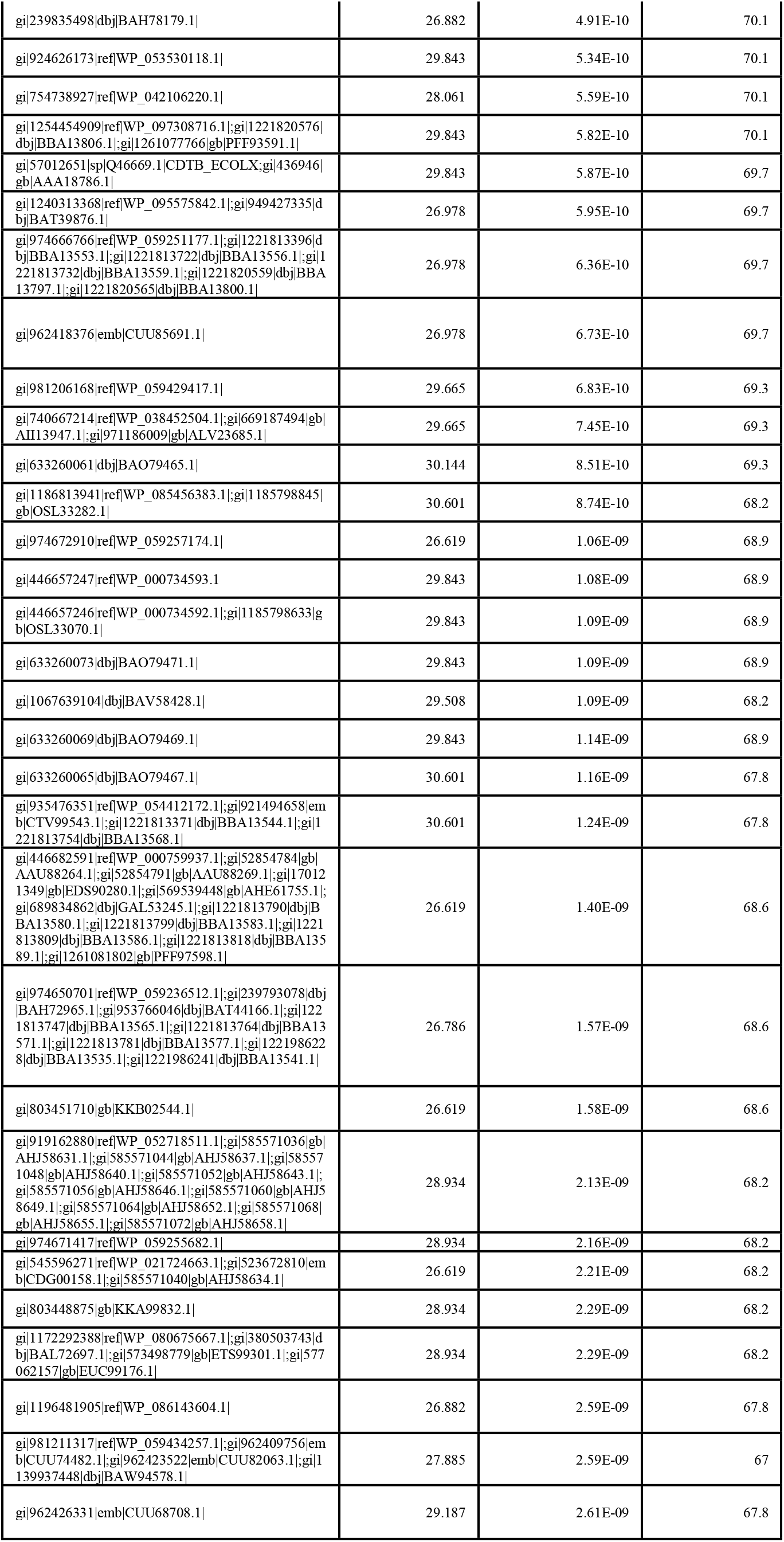
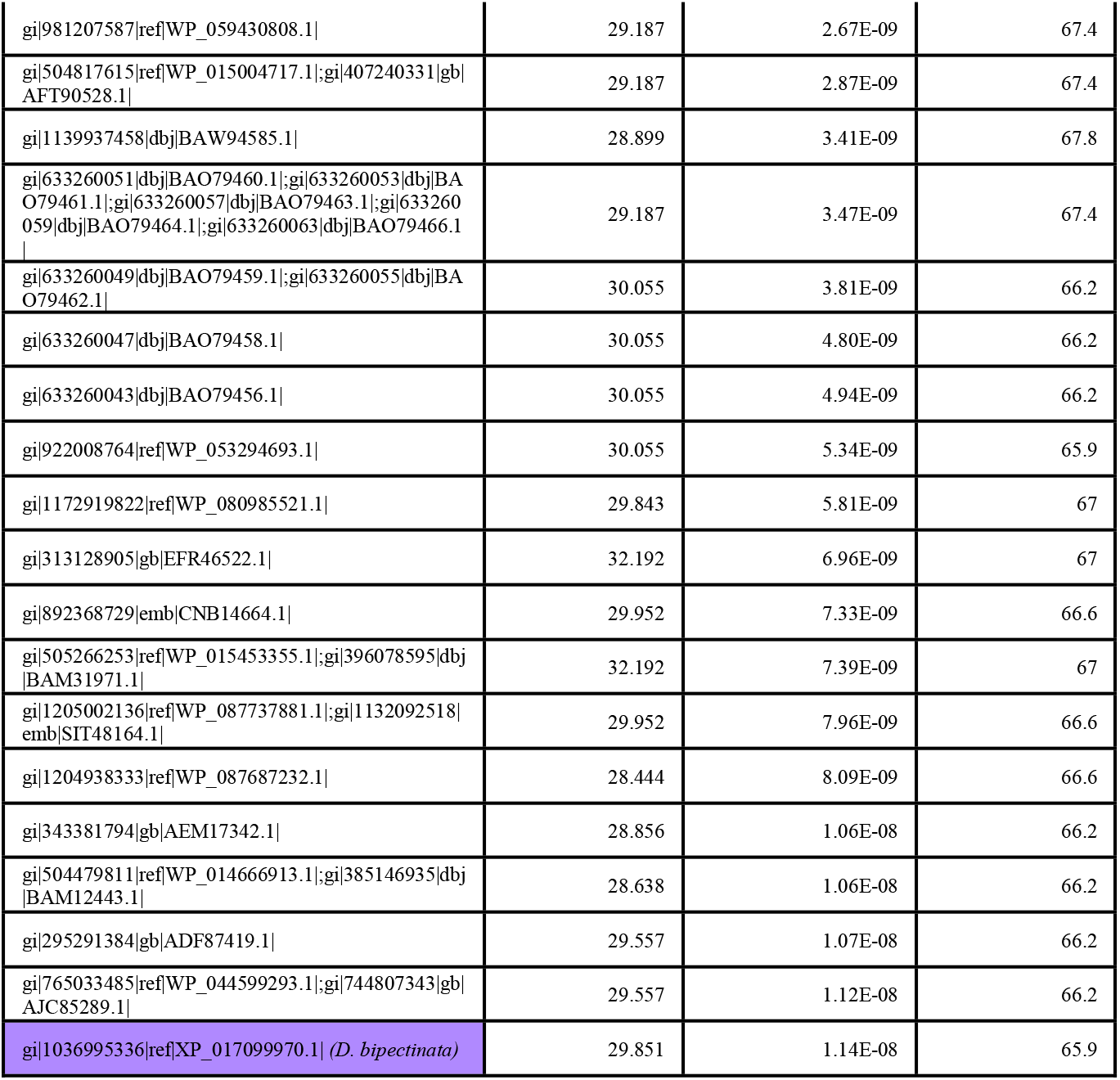
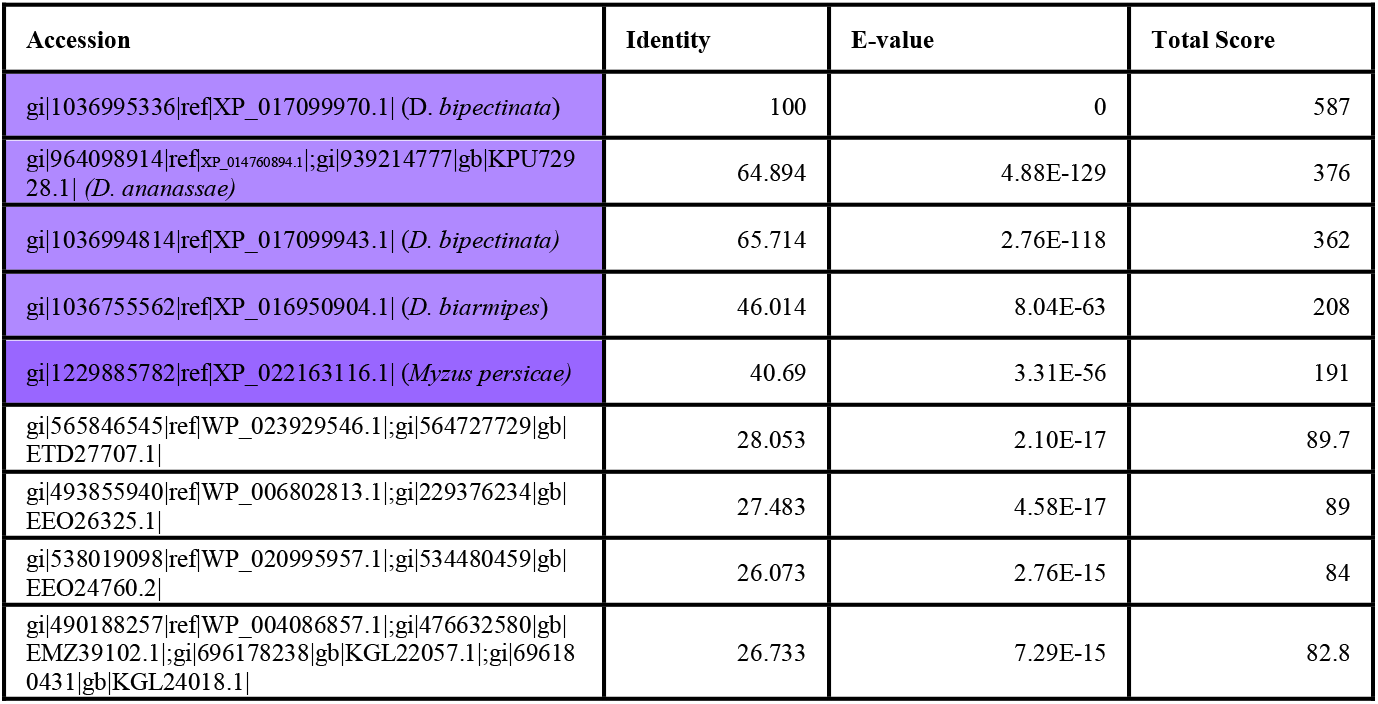
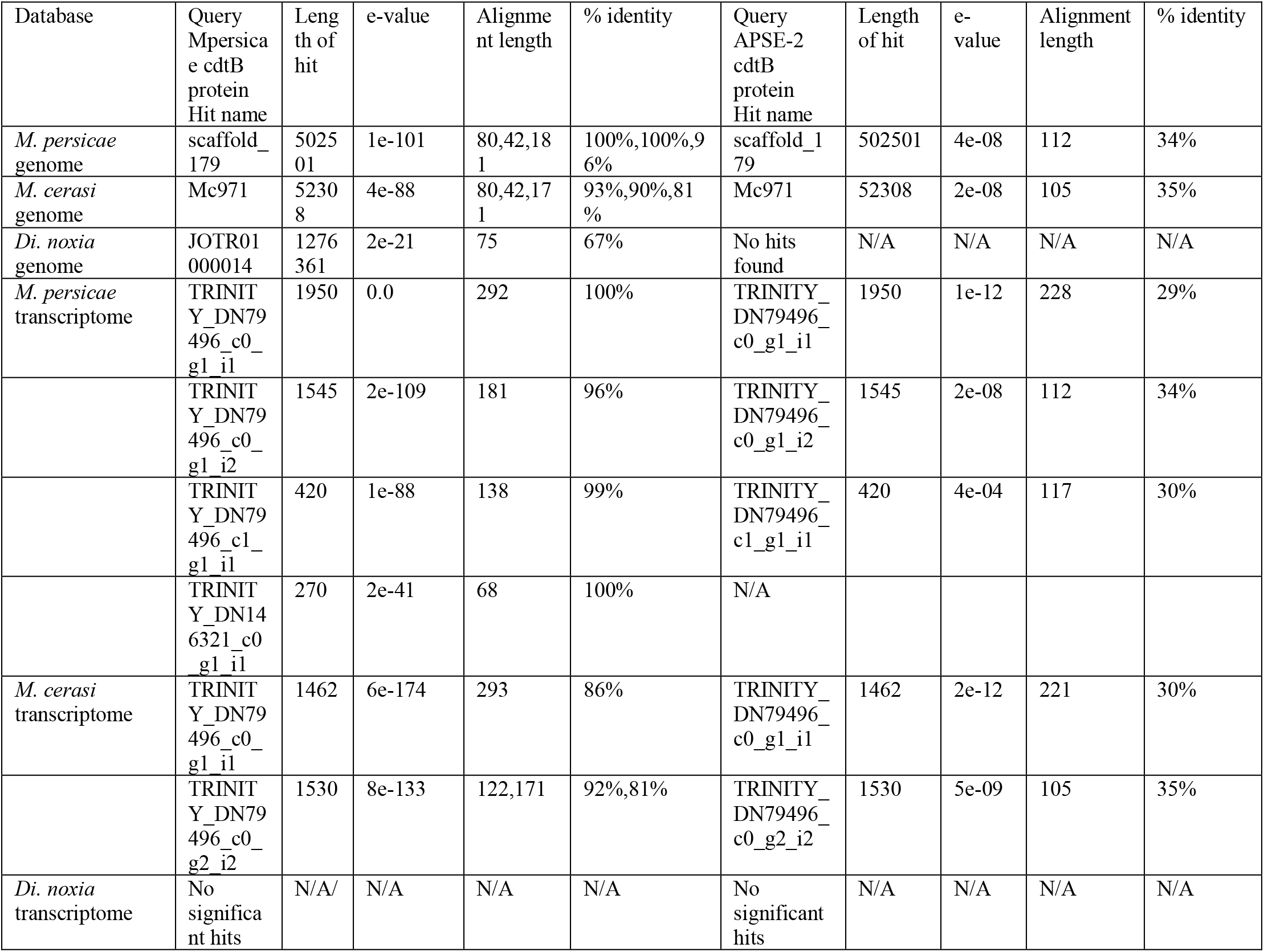
BLAST searches to the NCBI nr database were used to identify orthologs to *cdtB* from *S. flava*. For **Table S1a**, the TBLASTX search was run using default settings with *S. flava cdtB* as a query, which identified a *cdtB* ortholog in *Dr. bipectinata.* In **Table S1b**, BLASTP search was run using the *Dr. bipectinata* cdtB ortholog (NCBI ID: XP_017099970.1) as a query. Relevant matches are highlighted in purple with added description from authors in parentheses. **Table S1c** presents positive TBLASTN output results from searching *Myzus* genomes and transcriptomes using *M. persicae* cdtB (NCBI ID: XP_022163116) and *Ca.* H. defensa cdtB (UniProt ID: C4K6T7) as queries.

**Table S2.** Details of genome assemblies and scaffolds from representative drosophilids or aphids in which *cdtB* was identified.

|  | Genome | Genome Scaffold N50 | Genome Coverage | Scaffold Size | Exons |
| --- | --- | --- | --- | --- | --- |
| *Dr. ananassae* | GCA_000005115.1 | 4.59 Mb | 8.9X | 5.36 Mb (NW_001939298) | 3 (GF26441) |
| *Dr. bipectinata* | GCA_000233415.2 | 664 kb | 266.3X | 732 kb (KB464254.1) | 3 (LOC108127428) |
| *Dr. bipectinata* | GCA_000233415.2 | 664 kb | 266.3X | 732 kb (KB464254.1) | 5 (LOC108127405) |
| *Dr. biarmipes* | GCF_000233415.1 | 3.38 Mb | 186.9X | 3.95 Mb (KB462579.1) | 3 (LOC108025143) |
| *S. flava* | RKRM00000000.1 | 105 kb | 90X | 1.4 Mb | 1 |
| *Dr. primaeva* | Ellie Armstrong, unpublished | 272 kb | 71.5X | 207 kb | no transcriptome data available |
| *M. persicae* | GCF_001856785.1 | 435 kb | 200X | 502 kb | <sup>3</sup><br>(XM_022307424.1) |

**Table S3.**
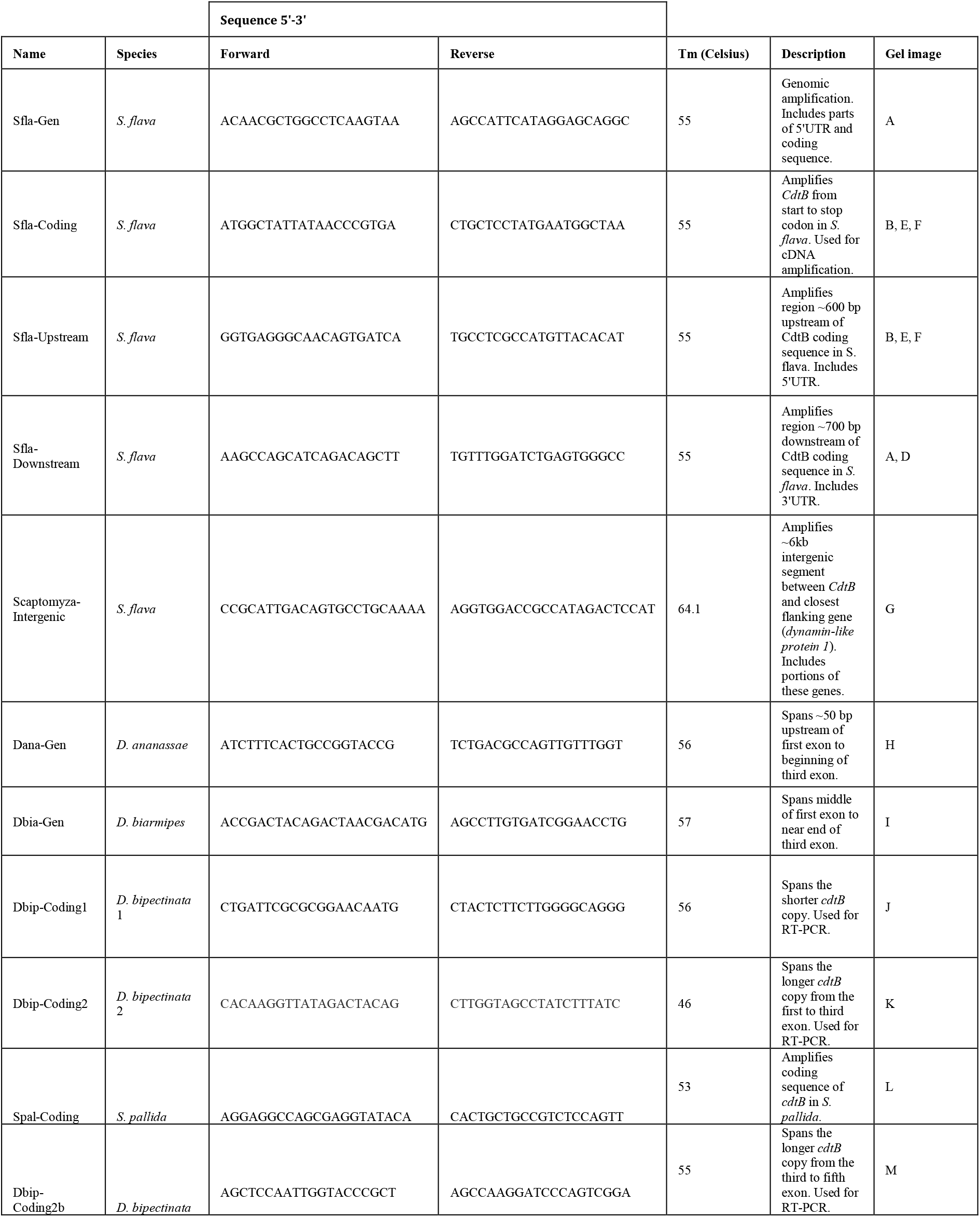
Primers used for PCR and RT-PCR. For associated gel images, please refer to **Figure S1;** for PCR conditions, please refer to **Materials and Methods.**

**Table S4.**
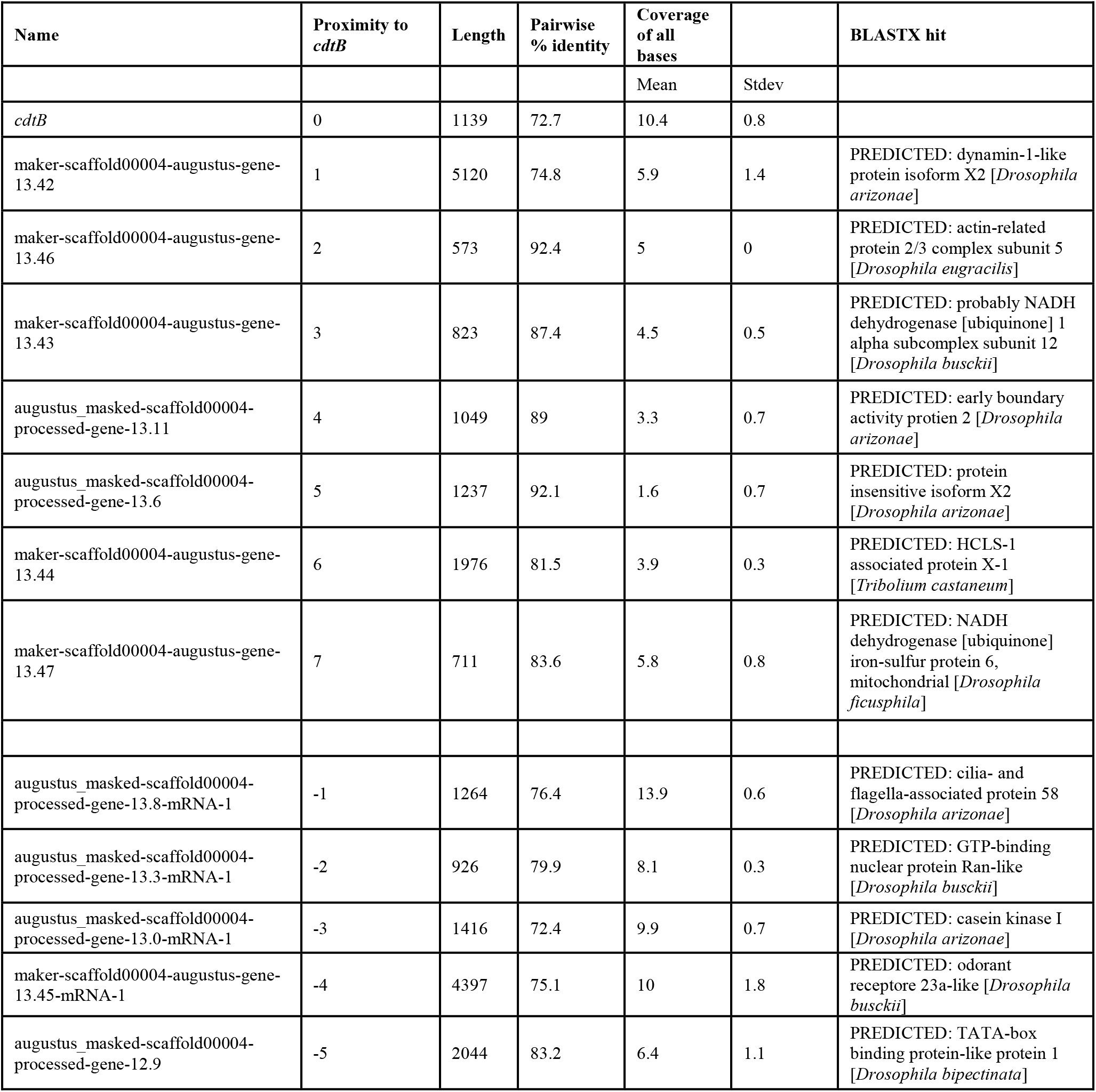
PacBio read alignment information for genes up and downstream of *cdtB* in *S. flava*. Gene homology was determined via highest-confidence BLASTX (*10*) hits (default settings) using genes predicted from the *S. flava* genome. ‘Proximity to *cdtB’* is negative if 5’ of *cdtB* and positive if 3’ of *cdtB*. For example, a value of −4 indicates that the gene is four predicted genes upstream (5’) of *cdtB.* For more information, please refer to the main text.

**Table S5.**
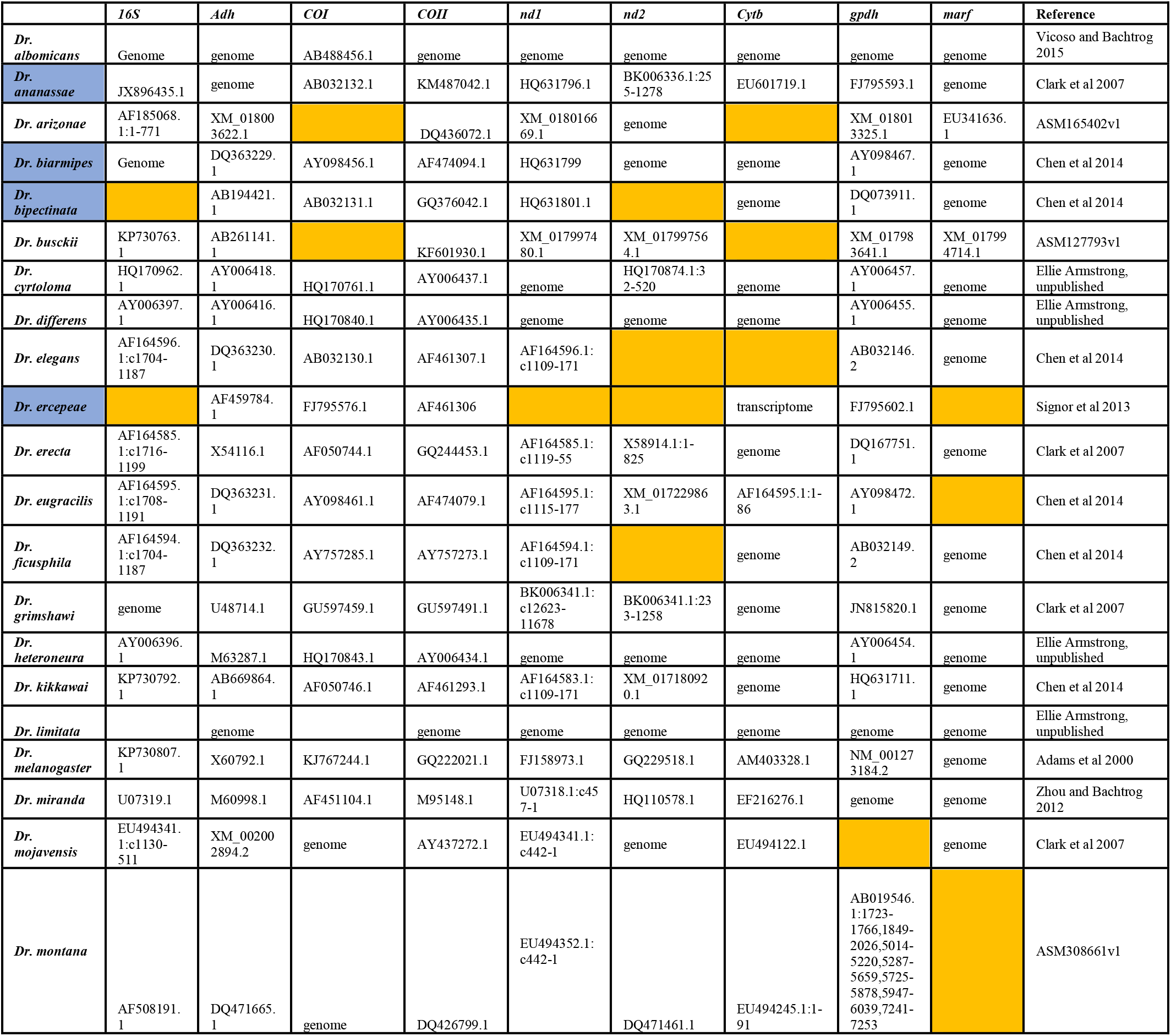
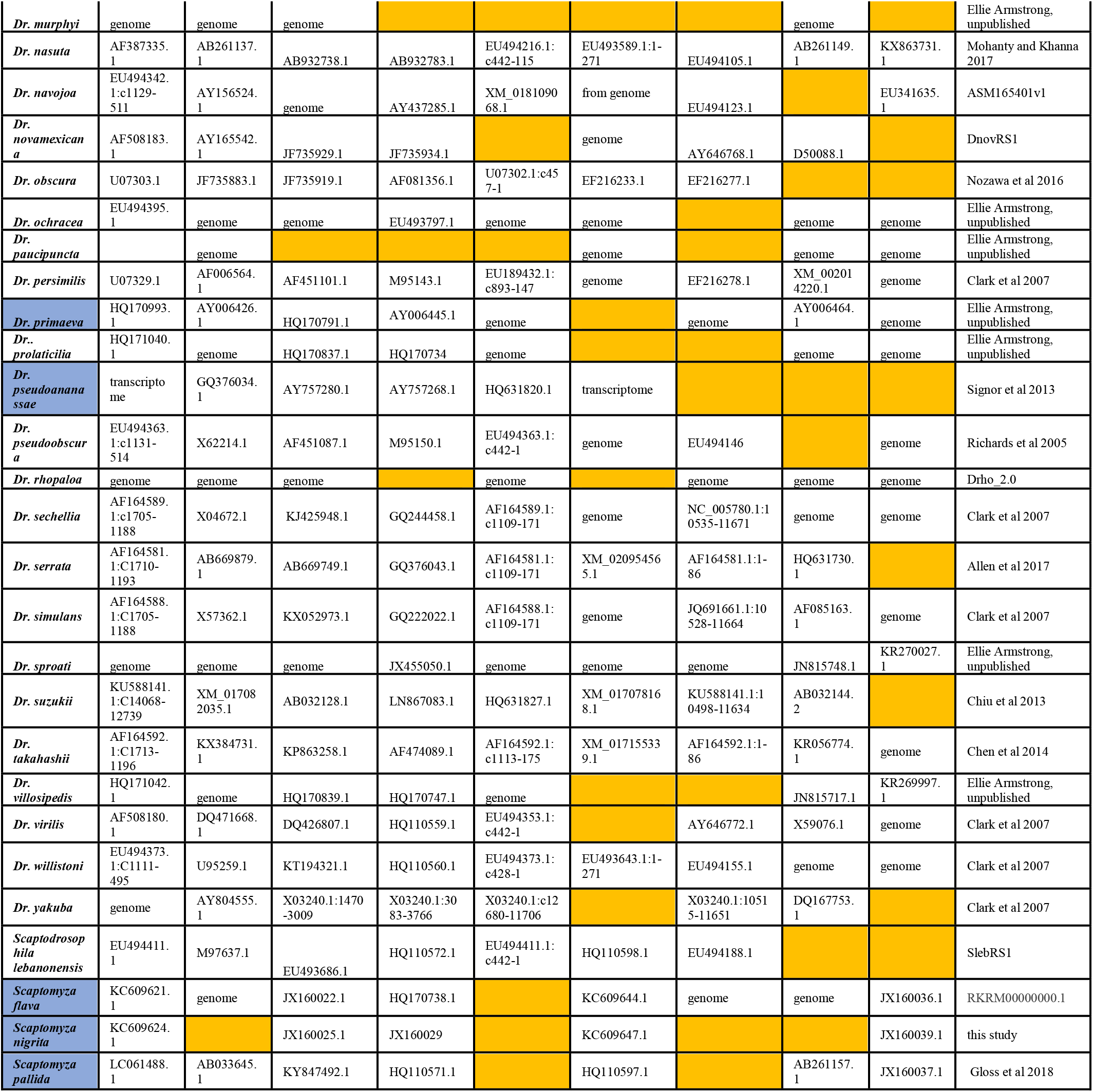
List of all drosophilid genomes (transcriptomes are included only if *cdtB* was found) screened for evidence of *cdtB.* Genomes or transcriptomes were searched using TBLASTX with default options using *S. flava, D. ananassae* and *Ca.* H. defensa *cdtB* as queries. A liberal E-value of 0.0001 was the cutoff to consider a positive identification of *cdtB.* Species in which *cdtB* was identified are highlighted in purple. Accession numbers of genes used to construct the drosophilid species phylogeny are included. If no sequences were found on GenBank, the genomes (cited in the column labeled ‘Reference’) were searched using BLASTN (using the *Dr. melanogaster* gene sequence as a query) and the highest-confidence hit was used; in this case, the corresponding cells are marked ‘genome’. If no sequences were found on GenBank or from searching the genome, the cell is highlighted yellow.

**Table S6.**
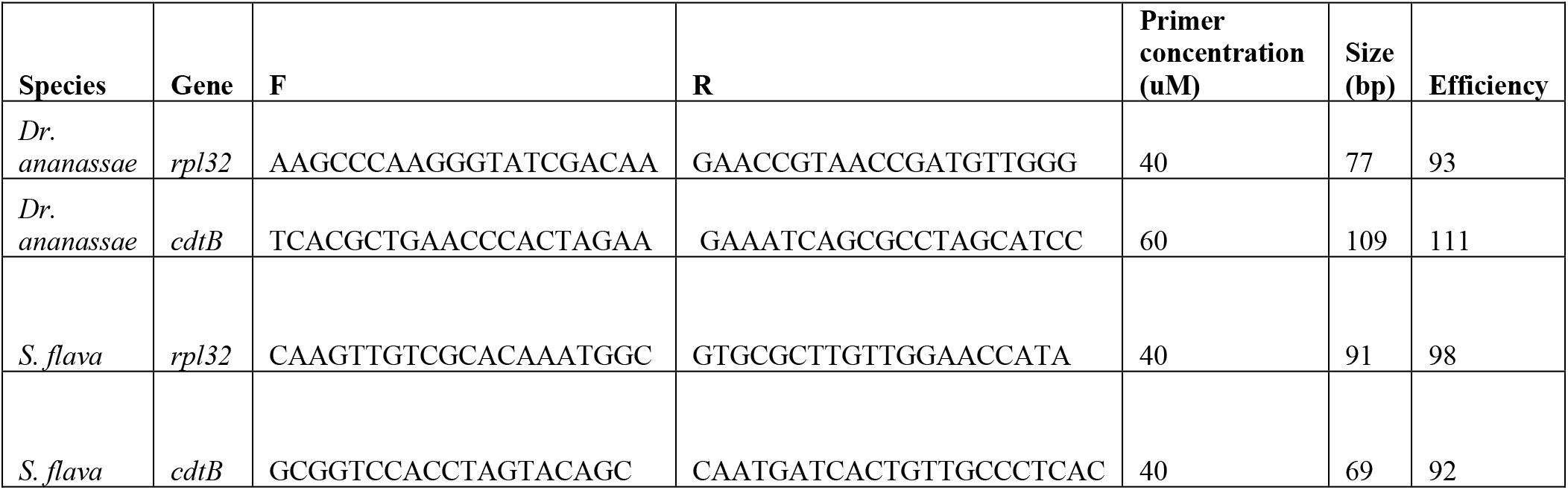
Primers used for RT-qPCR including concentrations used, size of the amplicon, and efficiencies. For description of reaction and cycling conditions please refer to **Methods**.

**Table S7.**
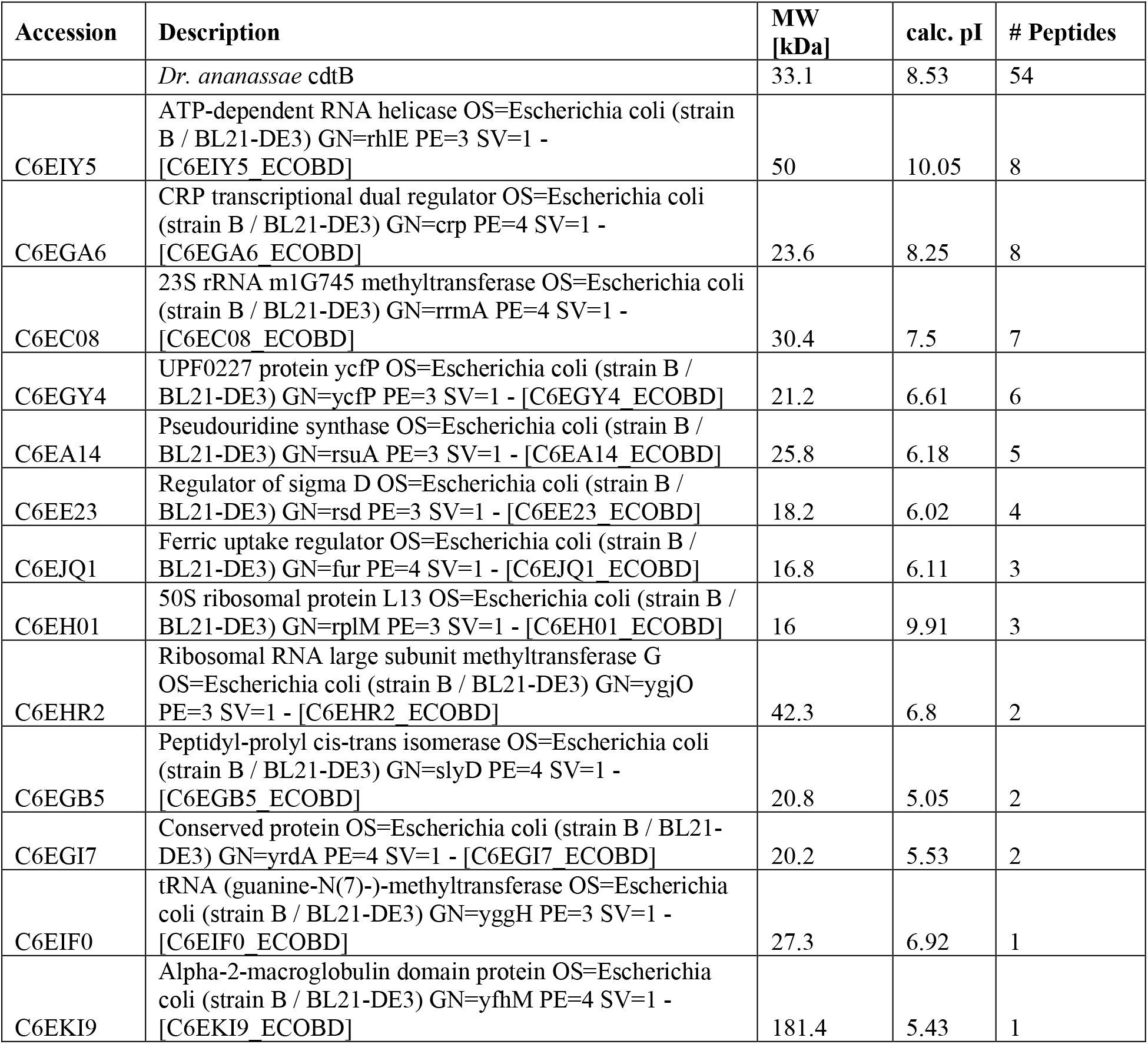
Mass spectrometry-based identification of the components of purified *Dr. ananassae* cdtB. ‘# peptides’ is equivalent to abundance of identified protein in purified *Dr. ananassae* cdtB solution.

**Table S8.**
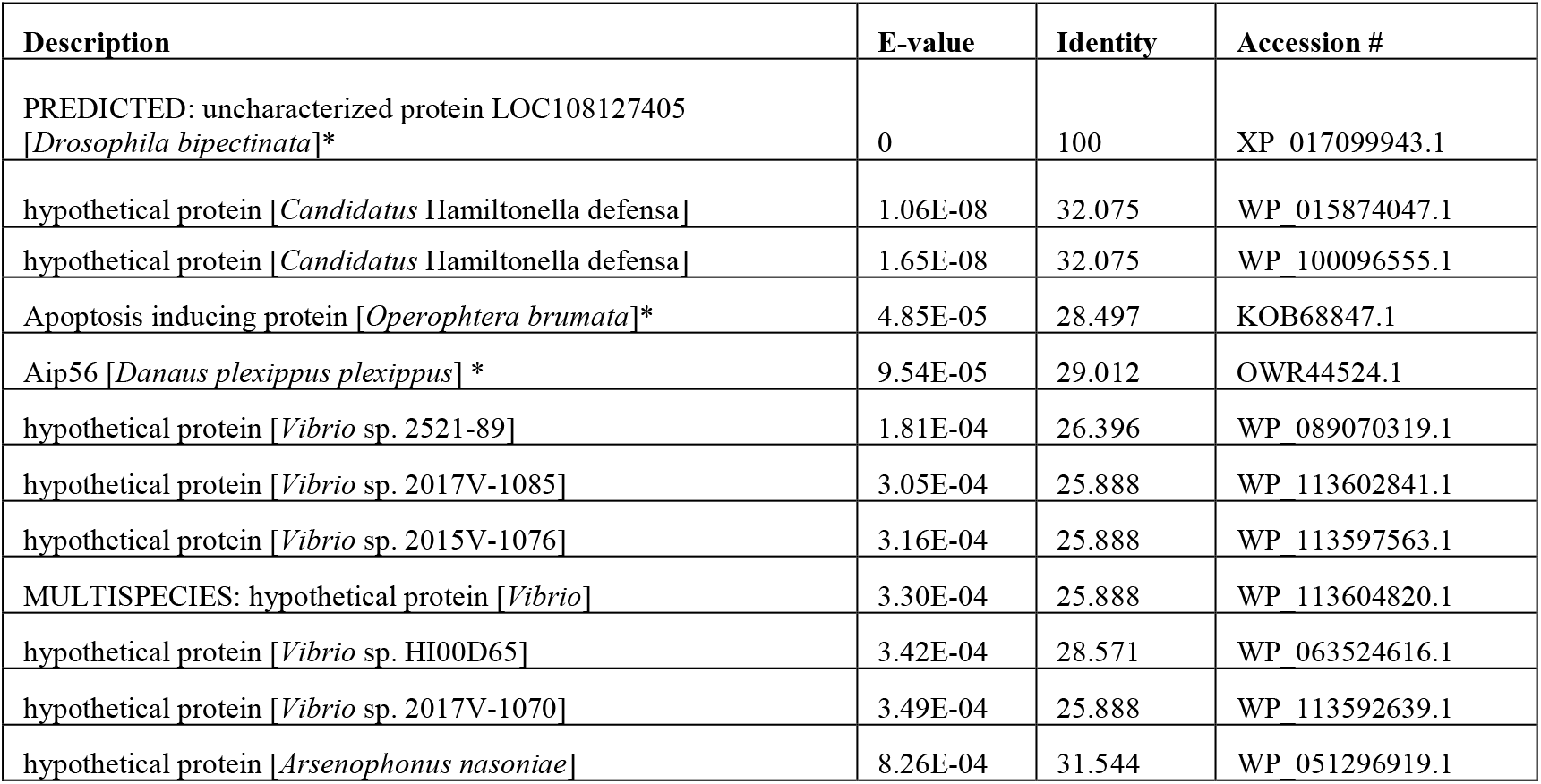
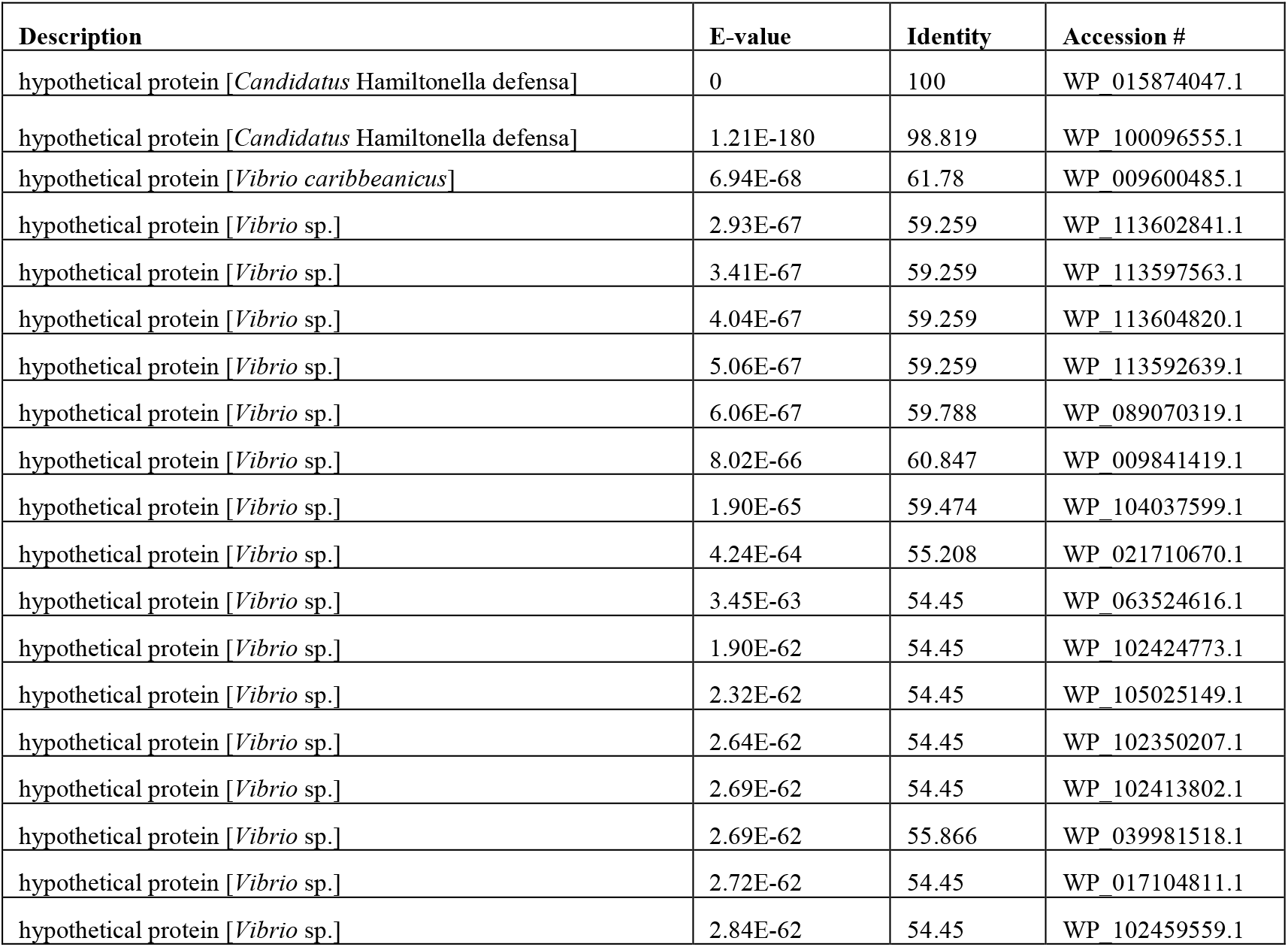
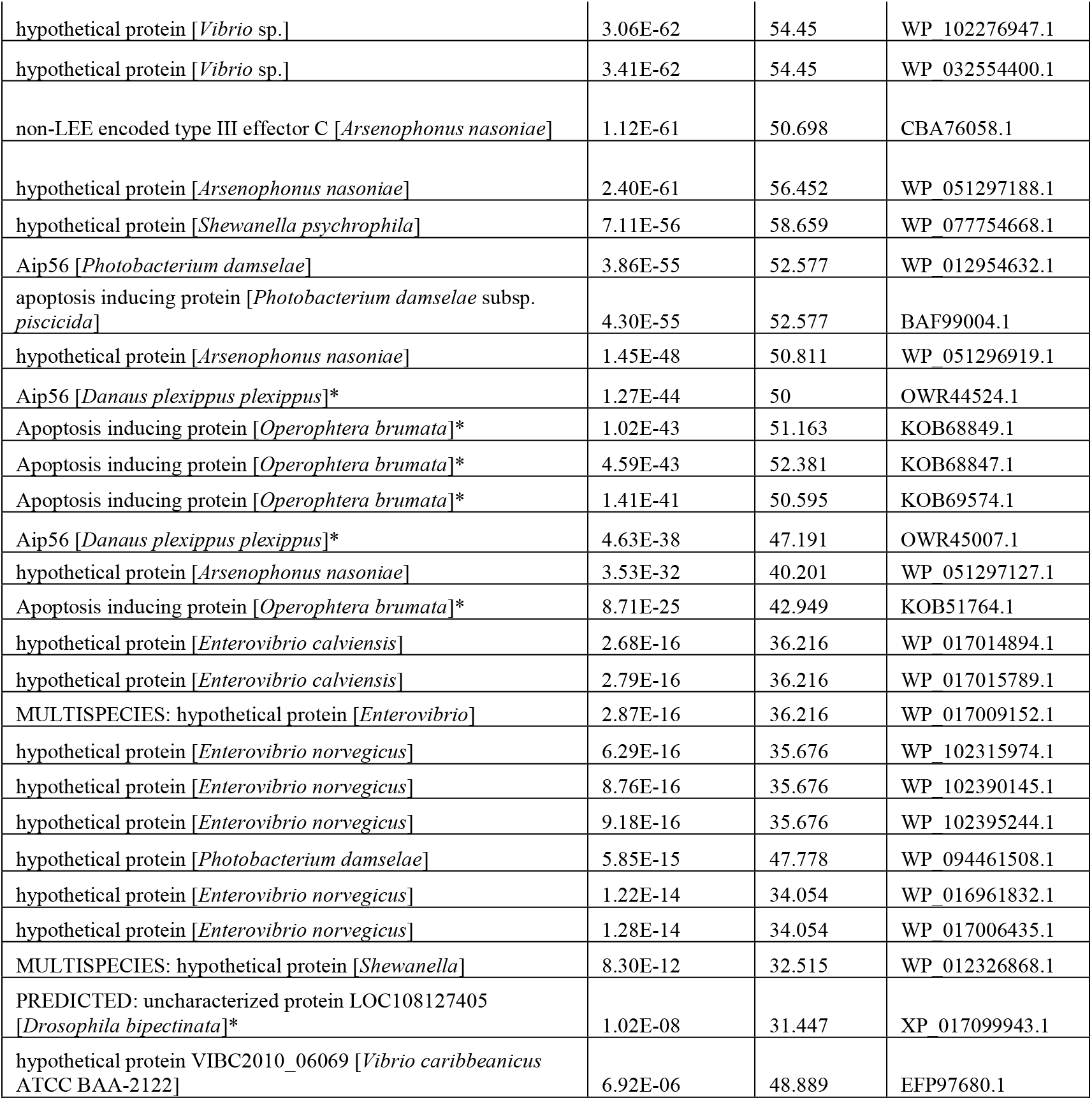
BLASTP results identifying the *Dr. bipectinata* cdtB + aip56 fusion. Results in eukaryotic species are indicated with an asterisk. **A.** BLASTP results from *Dr. bipectinata* non-alignable cdtB residues (residues 294-651 from XP_017099943.1) suggest homology to *apoptosis inducing protein 56 (aip56*). **B.** BLASTP results from *Ca.* H. defensa ORF D.

**Table S9.** Microsyntenic analysis of genes immediately up and downstream of *cdtB* in representative drosophilid species. Genes were determined in FlyBase (*Dr. ananassae*), from Chen et. al. 2014 (*85*) (*Dr. bipectinata* and *Dr. biarmipes*), the unpublished *S. flava* genome, or Mathers et. al. 2017 (*41*). Homology of up and downstream genes was predicted via default BLASTX searches using genes of interest as queries. ‘Proximity to *cdtB’* is described in **Table S4.** Synteny between the aphid species is supported by macrosyntenic analysis (see **Supplementary Text**).

|  | *D. ananassae* |  | *D. bipectinata* |  | *D. biarmipes* |  |
| --- | --- | --- | --- | --- | --- | --- |
| Proximity to *cdtB* | FlyBase ID | Predicted Homolog | Accession # | Predicted Homolog | Accession # | Predicted Homolog |
| -5 | Dana/GF22876 | **Mitotic checkpoint protein BUB3.** Has a dual function in spindle-assembly checkpoint signaling and in promoting the establishment of correct kinetochore-microtubule (K-MT) attachments. Promotes the formation of stable end-on bipolar attachments. | LOC108127480/XP_017100063.1 | **Mitotic checkpoint protein BUB3.** Has a dual function in spindle-assembly checkpoint signaling and in promoting the establishment of correct kinetochore-microtubule (K-MT) attachments. Promotes the formation of stable end-on bipolar attachments. | LOC108025083/XP_016950831.1 | **tRNA (adenine(58)-N(1))-methyltransferase catalytic subunit TRMT61A.** Belongs to family of transferases transferring one-carbon group methyltransferases. |
| -4 | Dana/GF22875 | **Lysoplasmalogenase-like protein TMEM86A.** Enzyme catalyzing the degradation of lysoplasmalogen, which are formed by the hydrolysis of the abundant membrane glycerophospholipids plasmalogens. May control respective level of plasmalogens and lysoplasmalogens in cells and modulate cell membrane properties. | LOC108127421/XP_017099965.1 | **Lysoplasmalogenase-like protein TMEM86A.** Enzyme catalyzing the degradation of lysoplasmalogen, which are formed by the hydrolysis of the abundant membrane glycerophospholipids plasmalogens. May control respective level of plasmalogens and lysoplasmalogens in cells and modulate cell membrane properties. | LOC108025084/XP_016950832.1 | **Uncharacterized protein.** Has predicted KIAA1430 superfamily domain. |
| -3 | Dana/GF23359 | **Head-specific guanylate cyclase.** May have a role in phototransduction. Molecular function: GTP binding, guanylate cyclase activity, heme binding, cGMP biosynthetic process, intracellular signal transduction, positive phototaxis, rhodopsin mediated signaling pathway, visual perception. | LOC108127469/XP_01700047.1 | **Head-specific guanylate cyclase.** May have a role in phototransduction. Molecular function: GTP binding, guanylate cyclase activity, heme binding, cGMP biosynthetic process, intracellular signal transduction, positive phototaxis, rhodopsin mediated signaling pathway, visual perception. | LOC108025050/XP_016950796.1 | **Neuronal PAS domain-containing protein 4.** Required for contextual memory in the hippocampus. |
| -2 | Dana/Obp99c | **Odorant binding protein.** | LOC10812740/XP_017100048.1 | **General odorant-binding protein 99a** | LOC108025437/XP_016951417.1 | **Suppressor protein SRP40.** Function not known. |
| -1 | Dana/GF233<br>61 | **Doublesex- and mab-3 related transcription factor A2.** Protein features are: DM DNA-binding domain. Molecular function is described by: sequence-specific DNA binding; transcription factor activity; sequence specific DNA binding. | LOC108127418/<br>XP_017099960.1 | **Doublesex- and mab-3 related transcription factor A2.** Protein features are: DM DNA-binding domain. Molecular function is described by: sequence-specific DNA binding; transcription factor activity; sequence specific DNA binding. | LOC108025555/XP_016951578.1 | **Uncharacterized protein.** Protein features include: FAD/NAD(P)-binding domain; glucose-methanol-choline oxidoreductase. |
| 0 | GF26441 | *cdtB* | LOC108127428 /<br>LOC108127405 | *cdtB* | LOC108025143 | *cdtB* |
| 1 | Dana/Obp99<br>d | **Odorant binding protein.** | LOC108127413/XP_017099952.1 | **Odorant-binding protein 99d** | LOC108025356/XP_016951305.1 | **Uncharacterized protein.** |
| 2 | Dana/Obp99<br>b-3 | **Odorant binding protein.** | LOC108127412/XP_017099951.1 | **General odorant-binding protein 99b-like** | LOC108025355/XP_016951303.1 | **Uncharacterized protein.** Contains conserved protein domain DUF3421, which is found in the fish toxin natterin and other uncharacterized proteins. |
| 3 | Dana/Obp99<br>b-1 | **Odorant binding protein.** | LOC108127419/XP_017099961.1 | **General odorant-binding protein 99b-like** | LOC108025354/XP_016951302.1 | **HGH1.** Predicted to be involved in ribosome biogenesis. |
| 4 | Dana/GF233<br>63 | **Calcium-binding protein P isoform X1.** | LOC108127415/XP017099958.1 | **Calcium-binding protein P isoform X2:** proteins that participate in calcium cell signaling pathways by binding to Ca<sup>++</sup> | LOC108025150/XP_016950914.1 | **Trypsin-1.** Member of the trypsin family of serine proteases. |
| 5 | Dana/GF233<br>64 | **Uncharacterized protein.** | LOC108127417/XP_017099959.1 | **Uncharacterized protein.** | LOC108025146/XP_016950911.1 | **Trypsin alpha.** |

| *S. flava* |  |  |
| --- | --- | --- |
| Genes up or downstream of *cdtB* | Annotation ID | Predicted Homolog |
| -5 | augustus_masked-scaffold00004-processed-gene-12.9 | **TATA-box binding protein-like protein 1.** Part of a specialized transcription system that mediates the transcription of most ribosomal proteins through 5'-TCT-3' motif. |
| -4 | maker-scaffold00004-augustus-gene-13.45 | **Odorant receptor 23a-like.** |
| -3 | augustus_masked-scaffold00004-processed-gene-13.0 | **Casein kinase I.** Serine/threonine selective enzymes that function as regulators of signal transduction pathways in most eukaryotes. |
| -2 | augustus_masked-scaffold00004-processed-gene-13.3 | **GTP-binding nuclear protein Ran-like.** GTP-binding protein involved in nucleocytoplasmic transport. Required for the import of protein into the nucleus and also for RNA export. |
| -1 | augustus_masked-scaffold00004-processed-gene-13.8 | **Cilia- and flagella-associated protein 58.** |
| 0 | No annotation ID | *cdtB* |
| 1 | maker-scaffold00004-augustus-gene-13.42 | **Dynamin-1-like protein.** Functions in mitochondrial and peroxisomal division. Required for normal rate of cytochrome c release and caspase activation during apoptosis. Required for formation of endocytic vesicles. |
| 2 | maker-scaffold00004-augustus-gene-13.46 | **Actin-related protein 2/3 complex subunit 5.** Component of Arp2/3 complex which is involved in regulation of actin polymerization and together with activating nucleation-promoting factor mediates formation of branched actin networks. |
| 3 | maker-scaffold00004-augustus-gene-13.43-mRNA-1 | **Probably NADH dehydrogenase [ubiquinone] 1 alpha subcomplex subunit 12.** Accessory subunit of the mitochondrial membrane respiratory chain NADH dehydrogenase. |
| 4 | Gloss et al 2018. / augustus_masked-scaffold00004-processed-gene-13.11 / [1387264-1388214] | **Early boundary activity protein 2.** Required for chromatin domain boundary function during early embryogenesis. |
| 5 | Gloss et al 2018. / augustus_masked-scaffold00004-processed-gene-13.6-mRNA-1 / [1389880,1391098] | **Protein insensitive isoform X2.** Can act as a transcriptional repressor and corepressor. |

